# Enhanced fitness of SARS-CoV-2 variant of concern B.1.1.7, but not B.1.351, in animal models

**DOI:** 10.1101/2021.06.28.450190

**Authors:** Lorenz Ulrich, Nico Joel Halwe, Adriano Taddeo, Nadine Ebert, Jacob Schön, Christelle Devisme, Bettina Salome Trüeb, Bernd Hoffmann, Manon Wider, Meriem Bekliz, Manel Essaidi-Laziosi, Marie Luisa Schmidt, Daniela Niemeyer, Victor Max Corman, Anna Kraft, Aurélie Godel, Laura Laloli, Jenna N. Kelly, Angele Breithaupt, Claudia Wylezich, Inês Berenguer Veiga, Mitra Gultom, Kenneth Adea, Benjamin Meyer, Christiane Eberhardt, Lisa Thomann, Monika Gsell-Albert, Fabien Labroussaa, Jörg Jores, Artur Summerfield, Christian Drosten, Isabella Anne Eckerle, Ronald Dijkman, Donata Hoffmann, Volker Thiel, Martin Beer, Charaf Benarafa

**Author notes:** These authors contributed equally to this work (Lorenz Ulrich, Nico Joel Halwe, Adriano Taddeo, Nadine Ebert). These authors jointly lead this work (Ronald Dijkman, Donata Hoffmann, Volker Thiel, Martin Beer, Charaf Benarafa).

## Abstract

Emerging variants of concern (VOCs) drive the SARS-CoV-2 pandemic. We assessed VOC B.1.1.7, now prevalent in several countries, and VOC B.1.351, representing the greatest threat to populations with immunity to the early SARS-CoV-2 progenitors. B.1.1.7 showed a clear fitness advantage over the progenitor variant (wt-S^614G^) in ferrets and two mouse models, where the substitutions in the spike glycoprotein were major drivers for fitness advantage. In the “superspreader” hamster model, B.1.1.7 and wt-S^614G^ had comparable fitness, whereas B.1.351 was outcompeted. The VOCs had similar replication kinetics as compared to wt-S^614G^ in human airway epithelial cultures. Our study highlights the importance of using multiple models for complete fitness characterization of VOCs and demonstrates adaptation of B.1.1.7 towards increased upper respiratory tract replication and enhanced transmission in vivo.

**Summary sentence:** B.1.1.7 VOC outcompetes progenitor SARS-CoV-2 in upper respiratory tract replication competition in vivo.

## Main Text

Uncontrolled transmission of severe acute respiratory syndrome-coronavirus 2 (SARS-CoV-2) in the human population has contributed to the persistence of the coronavirus disease 2019 (COVID-19) pandemic. The emergence of new variants in largely immunologically naïve populations suggests that adaptive mutations within the viral genome continue to improve the fitness of this zoonotic virus towards its host. In March 2020, the single amino acid change in the spike (S) protein at position 614 (S^614D^ to S^614G^) was identified in only a small fraction of sequenced samples but, within a few weeks, became the most predominant variant worldwide ^1^. The fitness advantage conferred by this single amino acid change was supported by a major increase in infectivity, viral loads, and transmissibility *in vitro* and in animal models ^2–4^.

In the second half of 2020, new SARS-CoV-2 variants of concern (VOCs) with a combination of several mutations emerged including the B.1.1.7 (also known as Alpha) first described in southeast England ^5^, and the B.1.351 (also known as Beta) first identified in South Africa ^6^. In spring 2021, B.1.1.7 rapidly became the prevailing variant in many regions of the world and a higher reproduction number was inferred from early epidemiological data ^7–9^. In addition to the S^614G^ change, B.1.1.7 has 18 different mutations in its genome, with two deletions and six substitutions within S alone ^10^. Some of the S mutations, such as N501Y and H69-V70del, were hypothesized to confer enhanced replication and transmission capability, but clear experimental evidence is lacking ^11, 12^. B.1.351 VOC has nine mutations in S, including N501Y and two in the S receptor-binding domain (RBD), K417N and E484K, the latter putatively responsible for escaping neutralization from convalescent patient plasma ^13–15^. Whether and which spike mutations are solely responsible for the putative fitness advantage is unknown.

Here, we investigated the fitness of the VOCs B.1.1.7 and B.1.351 compared to the predominant parental strain containing the S^D614G^ substitution – henceforth wt-S^614G^ – (i) in relevant primary airway culture systems *in vitro*, and (ii) in ferrets, Syrian hamsters, and two humanized mouse models to assess specific advantages in replication and transmission and to evaluate the impact of B.1.1.7 S mutations alone *in vivo*. Neither B.1.1.7 nor B.1.351 VOCs showed enhanced replication in human airway epithelial cell cultures compared to wt-S^614G^. Competitive transmission experiments in Syrian hamsters demonstrated similar replication and transmission of wt-S^614G^ and B.1.1.7, while both outcompeted B.1.351. However, competitive experiments in ferrets and transgenic mice expressing human angiotensin-converting enzyme 2 (hACE2) under the keratin 18 promoter (hACE2-K18Tg), which overexpress hACE2 in epithelial cells, showed increased fitness of B.1.1.7 compared to wt-S^614G^. Finally, B.1.1.7 and a recombinant clone expressing only the B.1.1.7-specific spike mutations (wt-S^B.1.1.7^) both outcompeted the parental wt-S^614G^ strain with higher virus load in the upper respiratory tract of knock-in mice (hACE2-KI), expressing solely hACE2 in place of mouse ACE2. In all *in vivo* models, B.1.1.7 and wt-S^614G^ infections resulted in similar pathology.

## Results

### Comparable replication for VOCs B.1.1.7, B.1.351, and wt-S^614G^ in primary nasal and bronchial airway epithelial cell cultures

To assess whether VOCs B.1.1.7 and B.1.351 exhibit a replicative advantage, we first infected primary nasal airway epithelial cell (AEC) cultures at an MOI of 0.02 at different ambient temperatures. No increased fitness was observed for clinical isolates of B.1.1.7 nor B.1.351 in comparison to a wild-type SARS-CoV-2 wt-S^614G^ clinical isolate (Fig. 1A,B). Because SARS-CoV-2 B.1.1.7 became the prevailing variant over progenitor variants (e.g., B.1.2) in many regions of the globe, we performed both individual replication and direct competition experiments to understand the potential replicative advantage conferred by mutations inside and outside the S region. Direct competition experiments in primary nasal and bronchial AEC cultures showed that clinical B.1.1.7 isolates and an isogenic variant with the B.1.1.7 spike (wt-S^B.1.1.7^) had replication kinetics similar to wt-S^614G^ when starting at equal titer ratios (Fig. 1C,D). Accordingly, individual B.1.1.7 spike mutations showed comparable replication kinetics to wt-S^614G^ (Extended Data Fig. 1). Plaque-reduction neutralization tests (PRNT) of plasma from convalescent (B.1.1.7 or non-VOC) patients and of serum from vaccinated (mRNA) individuals revealed no major difference in neutralization titers between wt-S^614G^ and isogenic recombinant variants expressing all or some S mutations of B.1.1.7 (Extended Data Fig 2A). PRNT of plasma samples from convalescent mild COVID-19 patients collected during the first pandemic wave showed lower neutralization titers of individual sera for B.1.1.7 and B.1.351 clinical isolates compared to the homologous B.1 (614G) isolate (Extended Data Fig 2B). Overall, the phenotype of both VOCs showed no major change in replication in human AEC cultures and were neutralized in part by serum of heterologous convalescent and vaccinated patients as reported in larger serological studies ^13–15^.

**Fig. 1.**
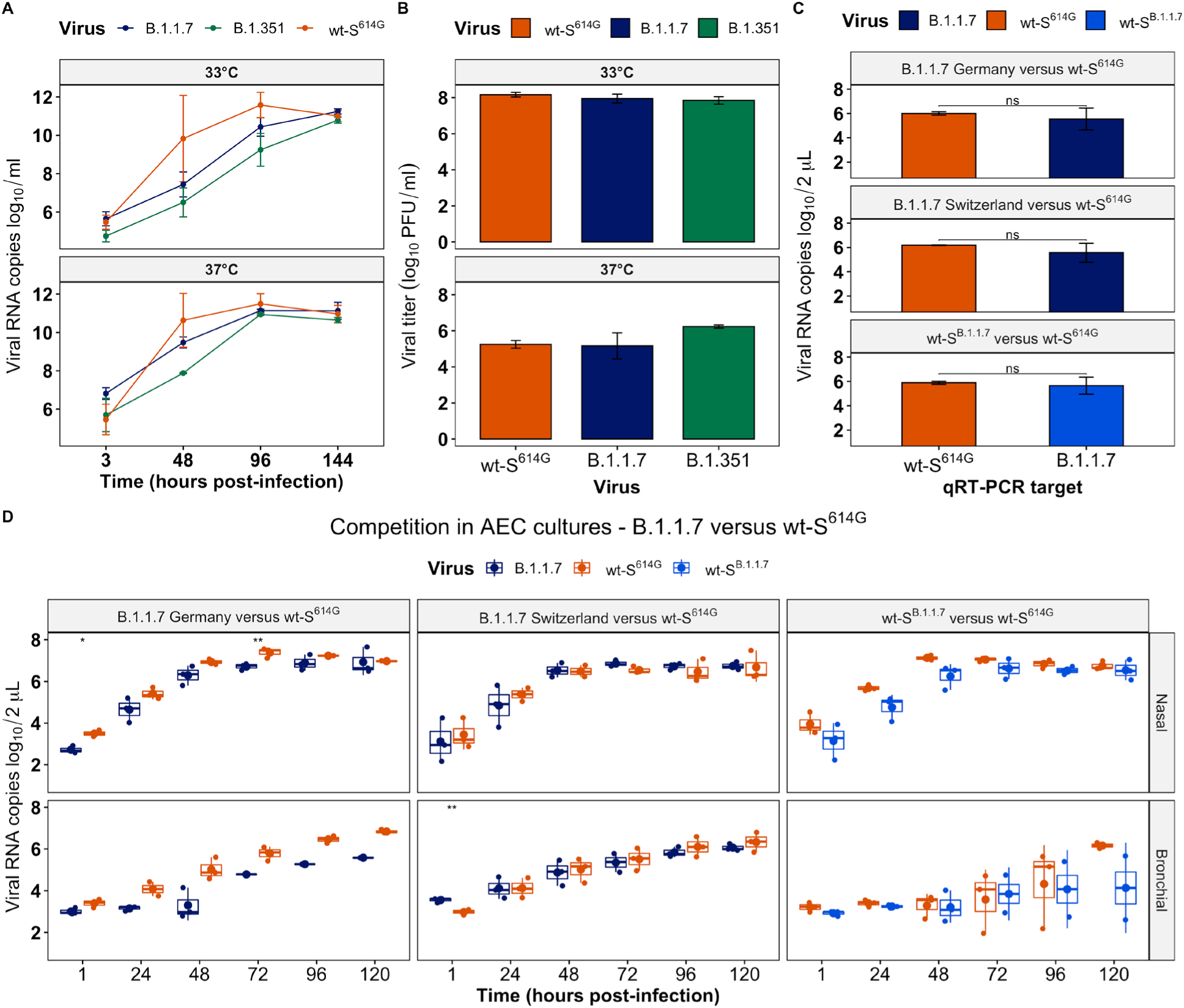
Replication kinetics of SARS-CoV-2 VOCs in primary airway epithelial cell cultures. **(A)** Viral replication kinetics of B.1.1.7, B.1.351, and wt-S^614G^ (MOI 0.02) at 33 and 37°C in primary airway epithelial cell (AEC) cultures derived from the nasal epithelium. **(B)** Quantification of corresponding infectious viral titers of B.1.1.7, B.1.351, and wt-S^614G^ 6 days post-infection. **(C)** Quantification of equal viral titer mixtures for SARS-CoV-2 B.1.1.7 and wt-S^B.1.1.7^ versus wt-S^614G^ used as inoculum to assess viral replication fitness in nasal and bronchial AEC cultures. **(D)** Corresponding viral kinetics during the direct competition assay of SARS-CoV-2 B.1.1.7 and wt-S^B.1.1.7^ versus wt-S^614G^ in primary nasal and bronchial AEC cultures. Data are presented as mean values ± standard deviations of two (A and B) or three (C and D) biological replicates, with error bars representing standard deviations. * p<0.05, ** p<0.01 by two-tailed t-test comparing B.1.1.7 and wt-S^614G^ at indicated time points (D).

### Comparable fitness for B.1.1.7 and wt-S^614G^, both outcompeting B.1.351 in hamsters

Groups of six Syrian hamsters were each inoculated intranasally with a mixture of two SARS-CoV-2 strains comprising comparable genome equivalents in three one-to-one competition experiments: B.1.1.7 *vs* B.1.351, B.1.351 *vs* wt-S^614G^, and B.1.1.7 *vs* wt-S^614G^. All experimentally infected “donor” hamsters were strictly kept in isolation cages to prevent intergroup spill-over infections. Each donor hamster was co-housed with a naïve “contact I” hamster 1 day post infection (dpi), creating six donor-contact I pairs to evaluate shedding and transmission (Extended Data Fig. 3A). On 4 dpi, donor hamsters were euthanized, and six subsequent transmission pairs were set up by co-housing each contact I hamster with a naïve contact II hamster (Extended Data Fig. 3A).

In two competition experiments, wt-S^614G^ and B.1.1.7 outcompeted B.1.351 in nasal washings of the donor hamsters from 1 dpi until euthanasia 4 dpi. Genome copies reached up to 10^9^ /mL for wt-S^614G^ and B.1.1.7, whereas B.1.351 viral loads were 10-fold lower at corresponding time points (Fig. 2, Extended Data Fig. 4). Consequently, transmission was dominated almost every time by the competing variants wt-S^614G^ (Fig. 2) and B.1.1.7 (Extended Data Fig. 4) in contact I and II hamsters, while B.1.351 was near the limit of detection in nasal washings (Fig. 2, Extended Data Figs. 4, 5B,C, 6C,D). In donor hamsters, viral genome loads in the upper respiratory tract (URT: nasal conchae and trachea) revealed higher replication of B.1.1.7 and wt-S^614G^ as compared to B.1.351 (Extended Data Figs. 7, 8). However, B.1.351 replicated to higher titers in the lower respiratory tract (LRT: cranial, medial, and caudal lung lobes) than B.1.1.7 and wt-S^614G^ in donor hamsters. Anyhow, a dominant transmission of B.1.1.7 and of S^614G^ compared to B.1.351 was observed with higher viral loads in the URT of contact I and II hamsters (Extended Data Figs. 7, 8).

**Fig. 2.**
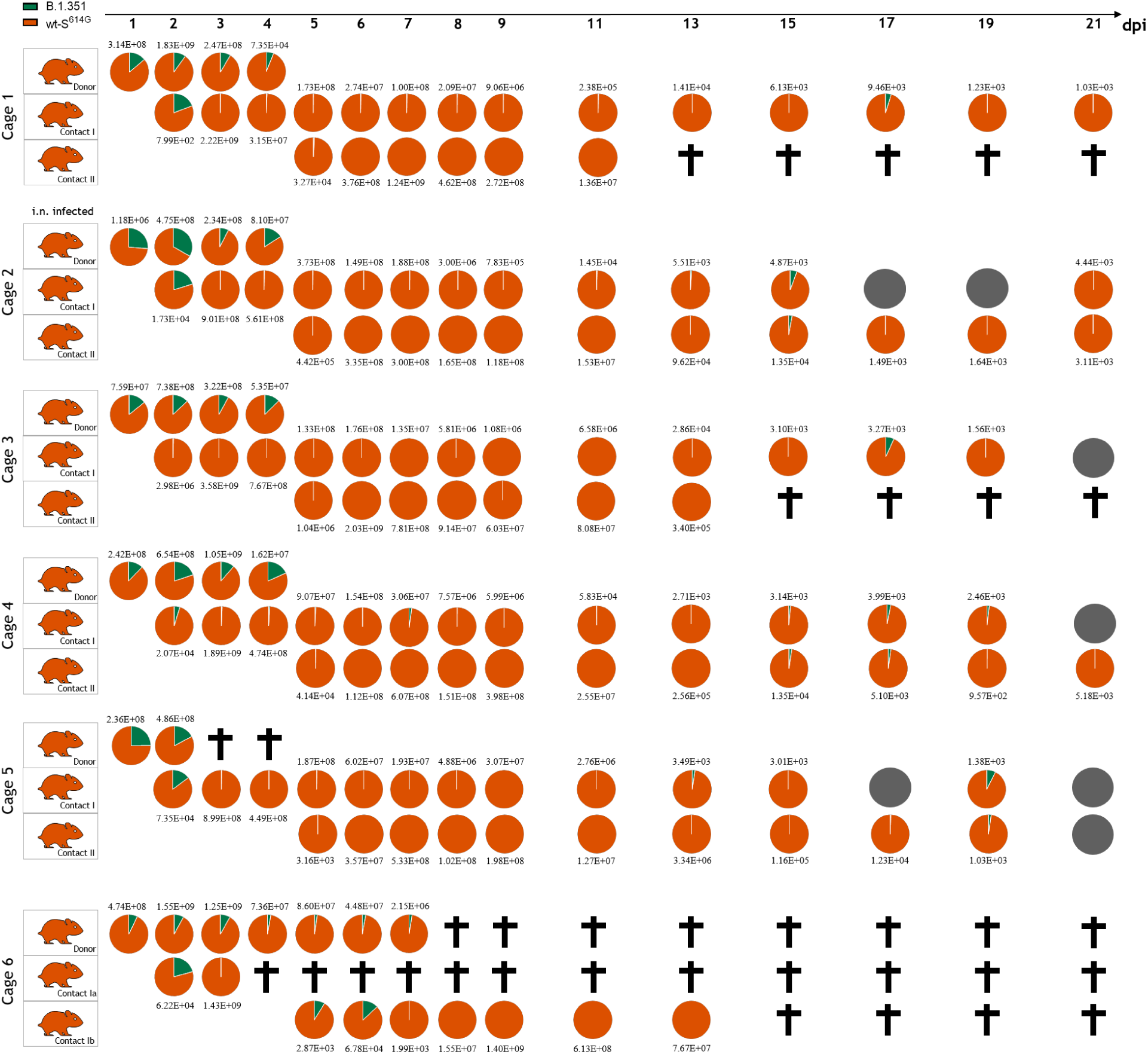
Competitive replication and transmission of B.1.351 and wt-S^614G^ in Syrian hamsters. Six donor hamsters were each inoculated with 10^4.25^ TCID_50_ determined by back titration and composed of a mixture of wt-S^614G^ (orange) and B.1.351 (green) at 1:3.8 ratio determined by RT-qPCR. Donor hamsters, contact I and II hamsters were co-housed sequentially as shown in Extended Data Fig. 3A. Nasal washings were performed daily from 1-9 dpi and afterwards every two days until 21 dpi. Each pie chart illustrates the ratio of the respective viruses in nasal washings for each sampling day. Grey pies indicate no detection of viral genomes. Total genome copies/ml are indicated above or below the respective pies. Hamster silhouettes are colored according to the dominant variant detected in the latest sample of each animal. Black crosses indicate the respective animal was already dead. Donor hamster in cage 5 and the contact I hamster of cage 6 passed away during inhalation anesthesia on 3 and 4 dpi, respectively, which required changes in the group composition where the donor hamster was kept until 7 dpi in the latter case and was co-housed in two different pairs: donor – contact Ia pair and another donor – contact Ib pair.

In the B.1.1.7 vs wt-S^614G^ competition, no clear fitness difference was observed in virus replication in nasal washes of donor hamsters, and both variants were detected at all time-points in each donor with B.1.1.7 specific copies ranging from 10^5^ to 10^9^ genome copies/mL, and wt-S^614G^ from 10^5^ to 10^8^ genome copies/mL (Fig. 3). Of note, B.1.1.7 was dominant over wt-S^614G^ in the donor hamsters at 1 dpi, but ratios were balanced by the endpoint at 4 dpi. In organ samples of the donor hamsters, the highest viral loads were confirmed in the LRT with at least 10-fold higher genome loads for B.1.1.7 in comparison to wt-S^614G^ for 5 out of 6 animals (Extended Data Fig. 9). Sequential transmission to contact I and II animals was highly efficient for both variants, which were simultaneously found in nasal washings of almost all contact hamsters, and equal dominance of B.1.1.7 and wt-S^614G^ in 3 out of 6 transmission groups (Fig 3, Extended Data Figs. 5A, 6A). No clear advantage for one variant over the other was noted in the viral loads of the URT and LRT at experimental endpoints (Extended Data Fig. 9).

**Fig. 3.**
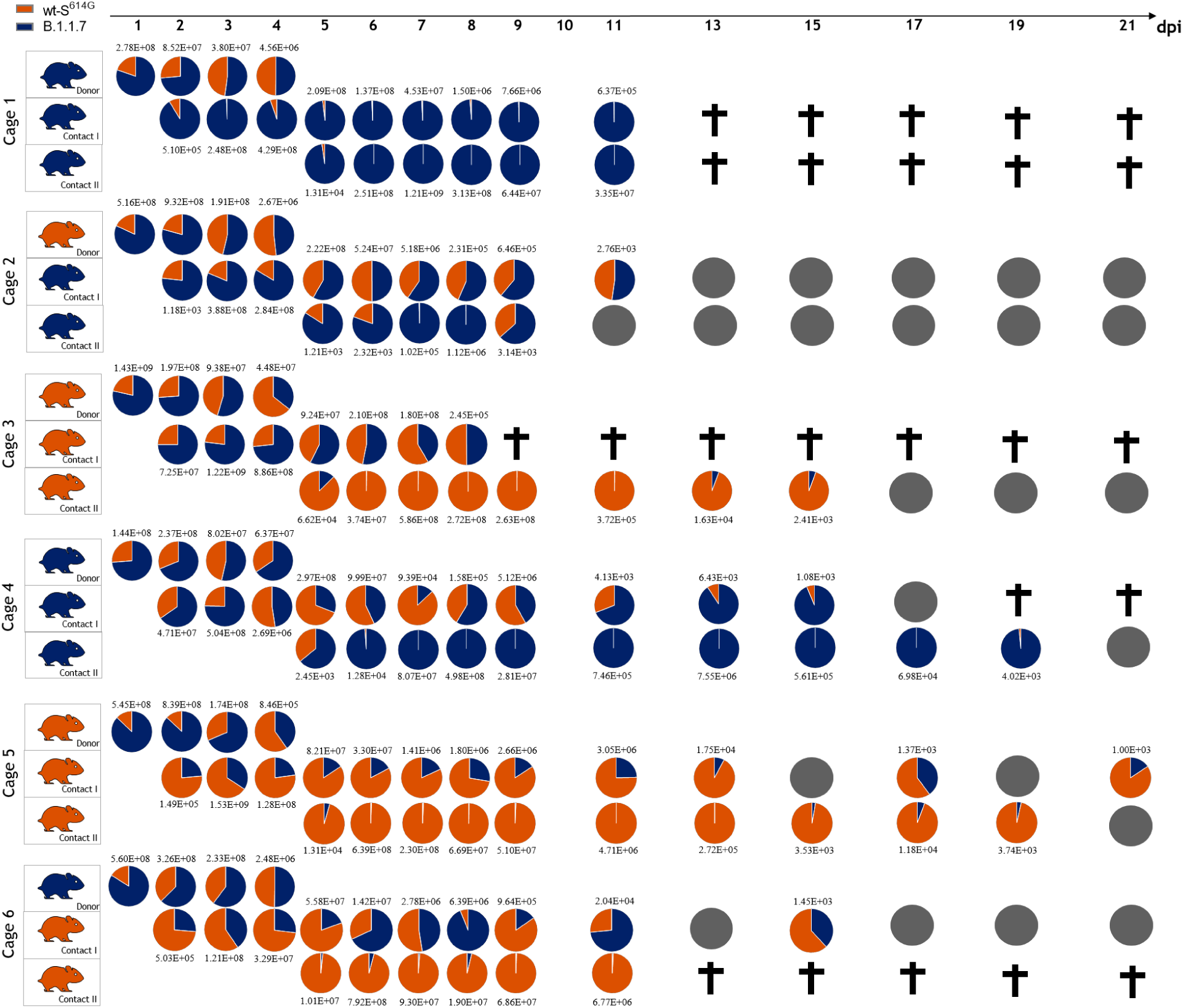
Competitive replication and transmission of B.1.1.7 and wt-S^614G^ in Syrian hamsters. Six donor hamsters were each inoculated with 10^4.3^ TCID_50_ determined by back titration and composed of a mixture of wt-S^614G^ (orange) and B.1.1.7 (dark blue) at 1:1.6 ratio determined by RT-qPCR. Donor hamsters, contact I and II hamsters were co-housed sequentially as shown in Extended Data Fig. 3A. Nasal washings were performed daily from 1-9 dpi and afterwards every two days until 21 dpi. Each pie chart illustrates the ratio of the respective viruses in nasal washings for each sampling day. Grey pies indicate no detection of viral genomes. Total genome copies/ml are indicated above or below respective pies. Hamster silhouettes are colored according to the dominant variant detected in the latest sample of each animal. Black crosses indicate that the corresponding animal reached the humane endpoint.

High SARS-CoV-2 replication in hamsters induced a rapid humoral immune response, as shown by serum reactivity in RBD-based ELISA and virus neutralization tests (VNT) (Extended Data Figs. 10, 11). Sera of all B.1.1.7 vs wt-S^614G^ contact I animals (20 days post co-housing with donors) were positive in both assays, while in the contact II group (17 days post co-housing with contact I) four out of six reacted positive in both tests, one negative only in the VNT and one negative in both assays. Reduced neutralization of B.1.351 by the sera of the hamsters of the three competition experiments is justified in part by the impaired replication and transmission of this virus and by the reduced cross-reactivity of the sera (Extended Data Figs. 10, 11).

### B.1.1.7 rapidly predominates over wt-S^614G^ in ferrets

In a similar approach as for hamsters, six donor ferrets were inoculated with a mixture of wt-S^614G^ and B.1.1.7 at comparable genome equivalents and sequential transmission was followed in naïve contact I and II ferrets (Extended Data Fig. 3B). B.1.1.7 rapidly became the dominant variant in nasal washings from 2 dpi with up to 10^5^ viral genome copies per mL (Fig. 4). Correspondingly, the nasal concha of donor ferrets revealed high replication in the nasal epithelium and up to 100- fold higher load of B.1.1.7 (up to 10^8.5^ viral genome copies per mL) than wt-S^614G^ (up to 10^6.5^ viral genome copies/ml) (Extended Data Fig. 12). While histopathological analysis clearly indicated viral replication within the nasal epithelium of the donor ferrets (Extended Data Fig. 13), no severe clinical signs were observed (Extended Data Figs. 5, 6). Transmission to contact I animals was only detected in two ferret pairs, from which only one contact I ferret transmitted the virus to the contact II ferret. However, in each of these three transmission events, B.1.1.7 variant was vastly dominant and replicated to similarly high titers as in donor ferrets (Fig. 4 and Extended Data Fig. 12). Virus shedding contact ferrets seroconverted by 21 dpi (Extended Data Figs. 10, 11).

**Fig. 4.**
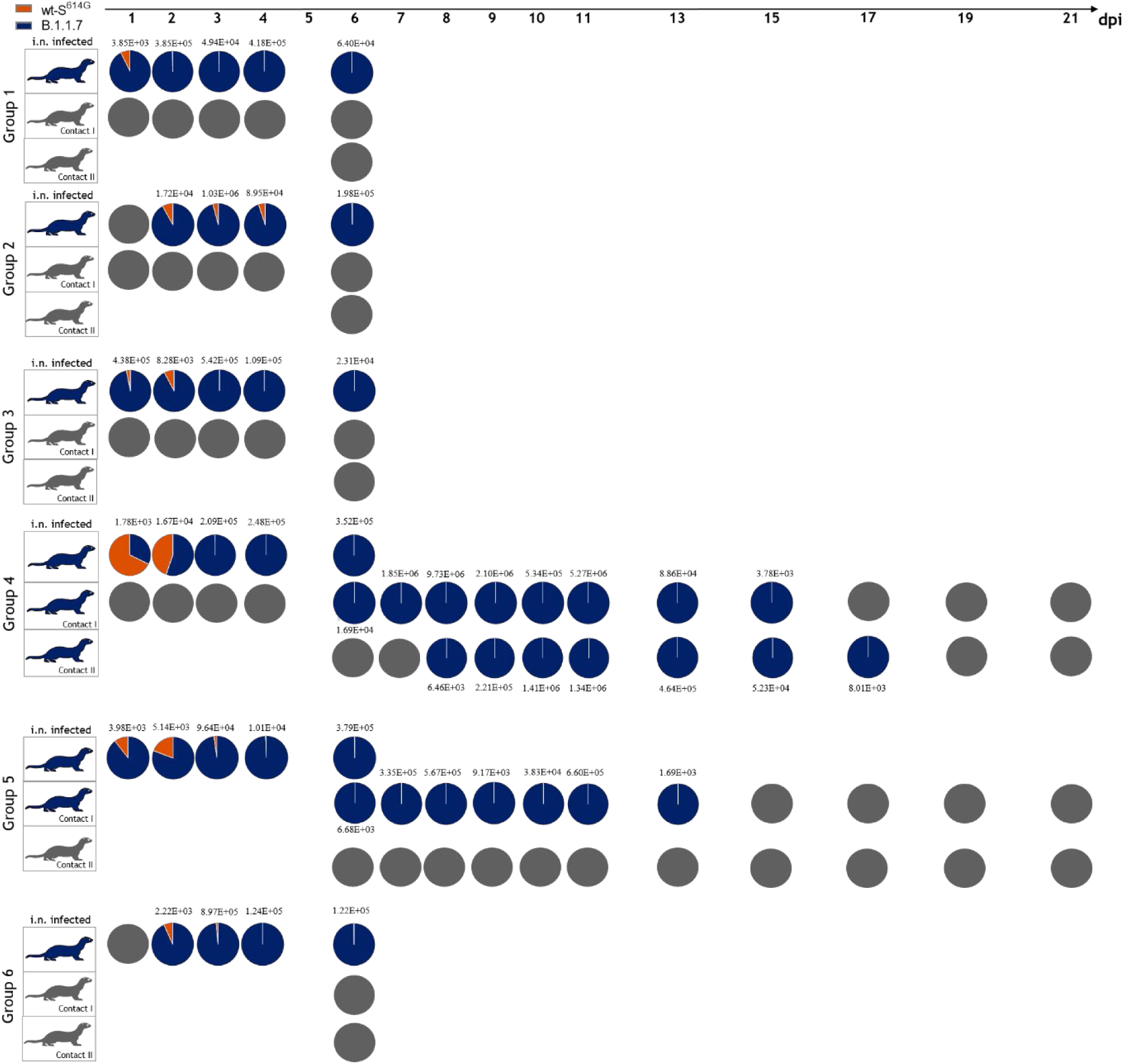
Replication and transmission of SARS-CoV-2 B.1.1.7 and wt-S^614G^ in ferrets. Six donor ferrets were each inoculated with 10^5.875^ TCID_50_ determined by back titration and composed of a mixture of wt-S^614G^ (orange) and B.1.1.7 (dark blue) at 1:1.2 ratio determined by RT-qPCR. Donor ferrets, contact I and II ferrets were co-housed sequentially as shown in Extended Data Fig. 3B. Pie charts illustrate the ratio of either SARS-CoV-2 B.1.1.7 or SARS-CoV-2 wt-S^614G^ detected in nasal washings of the donor or contact ferrets in the respective ferret groups at indicated dpi. Viral genome copies/mL are shown above or below respective pie charts; grey color indicates no detection of viral genomes. Coloring of the ferret silhouettes refers to the predominant SARS-CoV-2 variant detected in the latest sample of the respective animal.

### B.1.1.7 dominates the competition against wt-S^614G^ in hACE2-K18Tg transgenic mice

To assess further adaptation of B.1.1.7 to human ACE2, four hACE2-K18Tg mice, which overexpress hACE2 in respiratory epithelium ^16^, were inoculated with a mixture of SARS-CoV-2 wt-S^614G^ and B.1.1.7 at comparable genome equivalents (Fig. 5A). Each inoculated mouse was co-housed with a contact hACE2-K18Tg mouse at 1 dpi. A complete predominance of B.1.1.7 in the oropharyngeal samples of all four inoculated mice was observed from 1 to 4 dpi with up to 10^6^ viral genome copies/ml. This increased replicative fitness of B.1.1.7 over wt-S^614G^ was further reflected throughout the respiratory tract with higher genome copies in nose, lungs, olfactory bulb, and the brain at 4 dpi (Fig. 5A). All inoculated mice showed body weight loss at 4 dpi, but no weight loss was observed in contact mice (Extended Data Fig. 14A). Only B.1.1.7 viral genomes were detected in lungs of contact mice (Extended Data Fig. 14B).

**Fig. 5.**
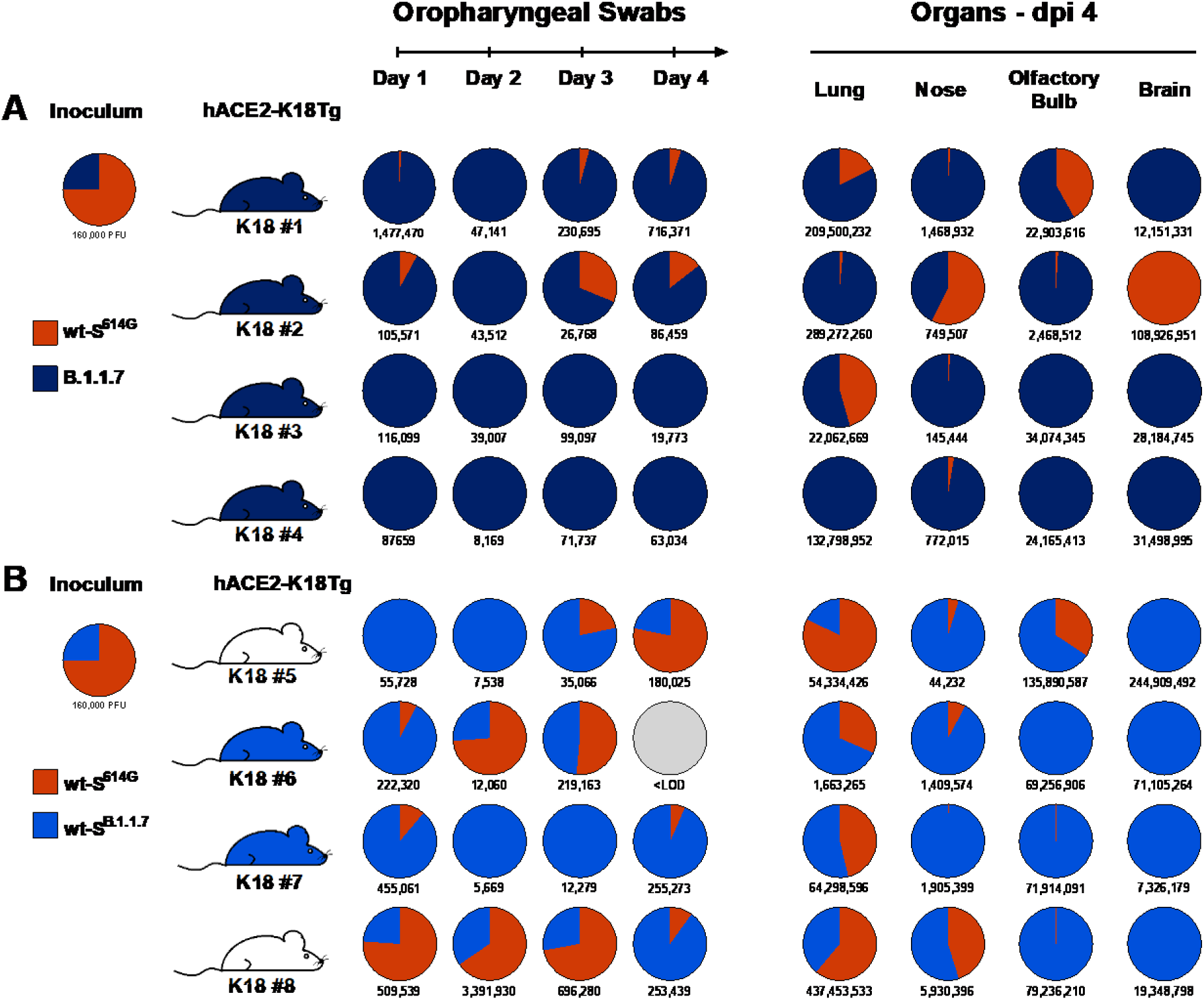
Replication of B.1.1.7, wt-S^B.1.1.7^, and wt-S^614G^ in hACE2-K18Tg mice. Two groups of four donor hACE2-K18Tg mice were inoculated with 1.6×10^5^ PFU per animal determined by back titration and composed of (**A**) a mixture of wt-S^614G^ (orange) and B.1.1.7 (dark blue) at 3:1 ratio determined by RT-qPCR, and (**B**) a mixture of wt-S^614G^ (orange) and wt-S^B.1.1.7^ (light blue) at 3:1 ratio determined by RT-qPCR. Pie charts illustrating the ratio of (**A**) wt-S^614G^ (orange) or B.1.1.7 (dark blue), and (**B**) wt-S^614G^ (orange) or wt-S^B.1.1.7^ (light blue) detected in the inoculum, oropharyngeal swabs, and tissues at the indicated day post inoculation (dpi). Total viral genome copies per mL are reported under the pie chart. Grey pies indicate samples under the limit of detection (<LOD, 10^3^ viral genome copies per organ). K18 #1 to #8 denote individual hACE-K18Tg donor mice. Coloring of the mouse silhouettes corresponds to the predominant SARS-CoV-2 variant, and white indicates absence of a clear dominance.

To differentiate the influence of the spike mutations from those of the rest of the genome of B.1.1.7, a similar competition experiment was performed between wt-S^614G^ and wt-S^B.1.1.7^. Transgenic hACE2-K18Tg mice were inoculated with a mixture of wt-S^B.1.1.7^ and wt-S^614G^ at comparable genome equivalents and placed with a contact hACE2-K18Tg mouse at 1 dpi. Interestingly, the replicative advantage of wt-S^B.1.1.7^ was less clear, especially in the lungs, where both wt-S^B.1.1.7^ and wt-S^614G^ variants showed comparable RNA levels (Fig. 4B). While none of the contact mice lost weight, two of the four mice had detectable viral RNA in the lung 7 days after contact and only wt-S^B.1.1.7^ was detected (Extended Data Fig. 14). The results indicate that the S^B.1.1.7^ spike mutations contribute partially to the replication advantage in the URT and to enhanced transmission of B.1.1.7 using mice that express high levels of human ACE2 in a K18-dependent manner.

### B.1.1.7 dominates wt-S^614G^ in the upper respiratory tract of hACE2-KI mice

We next used hACE2-KI homozygous mice, which express hACE2 in place of mouse ACE2 under the endogenous mouse *Ace2* promoter ^4^. In contrast to hACE2-K18Tg mice, hACE2-KI mice have a physiological expression of hACE2 with no ectopic expression of hACE2 in the brain, and no expression of mouse Ace2, which has been shown to be permissive to the spike mutation N501Y contained in S^B.1.1.7^ ^17^. Three groups of hACE2-KI mice were inoculated intranasally with a relatively low dose (10^4^ PFU/mouse) of either SARS-CoV-2 wt-S^614G^, B.1.1.7, or wt-S^B.1.1.7^ (n=8/group) as single virus infection (Fig. 6A). Significantly higher viral genome copy numbers were observed in mice infected with B.1.1.7 or with wt-S^B.1.1.7^ compared to wt-S^614G^ in oropharyngeal swabs 1 dpi, in the nose 2 dpi, and olfactory bulb 4 dpi (Fig. 6B,C). Of note, virus titers in the nose and lungs showed SARS-CoV-2 persistence 4 dpi in 3 out of 4 mice infected either with B.1.1.7 or with wt-S^B.1.1.7^, but not for hACE2-KI mice inoculated with wt-S^614G^ (Fig. 6D). No difference in lung histopathology score was observed between groups (SI Table 1). Finally, we performed competition experiments to compare the replication of the B.1.1.7 or wt-S^B.1.1.7^ with wt-S^614G^ in two groups of six hACE2-KI mice. We observed a complete predominance of B.1.1.7 variant and wt-S^B.1.1.7^ over wt-S^614G^ from 1 dpi in the URT. At 4 dpi, only B.1.1.7 or wt-S^B.1.1.7^ was detectable in lungs or nose (Fig. 5E,F). No weight loss was observed in any of the hACE2-KI mice (Extended Data Fig. 15). Together, the two humanized mouse models support enhanced fitness of SARS-CoV-2 B.1.1.7 variant over its ancestor wt-S^614G^, with increased replication and persistence in the URT and better systemic spread, mediated in part by changes located within the B.1.1.7 spike sequence. In binding assays in vitro, affinity between S^B.1.1.7^ and the ACE2 receptor was consistently higher than that of S^614G^ and ACE2 of humans, hamsters, and ferrets (Extended Data Fig. 16).

**Fig. 6.**
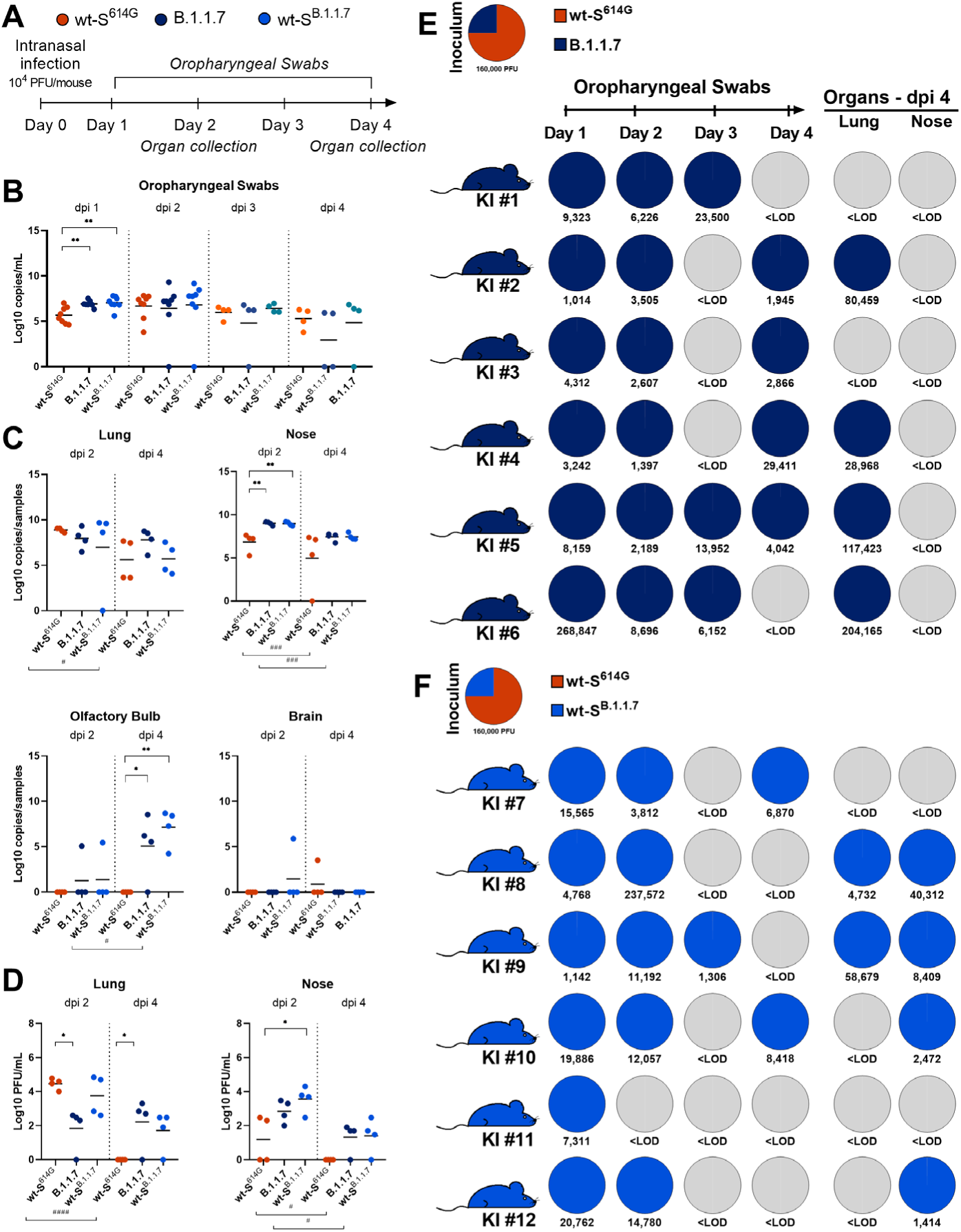
Replication of B.1.1.7 and wt-S^B.1.1.7^ in single virus and competition infections in hACE2-KI mice. (**A**) Experimental scheme for intranasal inoculation of groups of eight hACE2-KI mice with 10^4^ PFU per mouse of wt-S^614G^ (orange), or B.1.1.7 (dark blue) or wt-S^B.1.1.7^ (light blue). (**B, C**) Viral genome copy numbers in daily oropharyngeal swabs (**B**), and in tissues (**C**) at the indicated dpi. (**D**) Virus titers in tissues. Data were log10 transformed and presented as mean and individual values. * p<0.05, **p<0.01 by one-way ANOVA with Tukey’s multiple comparisons test comparing the three groups. # p<0.05, ### p<0.001 by two-tailed t-test comparing two- and four-days post inoculation. (**E, F**) Two groups of six donor hACE2-KI mice were inoculated with 1.6×10^5^ PFU determined by back titration and composed of (**E**) a mixture of wt-S^614G^ (orange) and B.1.1.7 (dark blue) at 3:1 ratio determined by RT-qPCR, and (**F**) a mixture of wt-S^614G^ (orange) and wt-S^B.1.1.7^ (light blue) at 3:1 ratio determined by RT-qPCR. Pie charts illustrating the ratio of (**E**) wt-S^614G^ (orange) or B.1.1.7 (dark blue), and (**F**) wt-S^614G^ (orange) or wt-S^B.1.1.7^ (light blue) measured in the inoculum, oropharyngeal swabs, and tissues at the indicated dpi. Total viral genome copies per mL are reported under the pie chart. Grey pies indicate samples under the limit of detection (<LOD, 10^3^ viral genome copies per organ). KI #1 to #12 denote individual hACE2-KI mice. Coloring of the mouse silhouettes refers to the corresponding predominant SARS-CoV-2 variant.

## Discussion

### Increased fitness of VOC B.1.1.7 in ferrets and humanized ACE2 mouse models

Epidemiological data indicate that new SARS-CoV-2 variant lineages with specific amino acid changes have a fitness advantage over contemporary strains. VOCs such as B.1.1.7 and B.1.351 are particularly concerning for their hypothesized ability to supersede progenitor strains and immune escape properties, respectively. In this comprehensive study, we provide experimental evidence that SARS-CoV-2 B.1.1.7 has a clear replication advantage over wt-S^614G^ in both ferret and two humanized mouse models. Moreover, B.1.1.7 exclusively was transmitted to contact animals in competition experiments, where ferrets and hACE2-K18Tg mice were inoculated with mixtures of B.1.1.7 and wt-S^614G^. We have also shown that the molecular mechanism behind the fitness advantage of B.1.1.7 *in vivo* is largely dependent on a few changes in the S including three amino acid deletions (H69, V70, Y144), and six substitutions (N501Y, A570D, P681H, T716I, S982A, D1118H). *In vitro*, B.1.1.7 spike mutations increased the affinity to human ACE2 over 5-fold, and 2-3-fold to ferret and hamster ACE2 indicative of an overall improvement in binding abilities rather than a specialization to human ACE2. In hACE2-KI mice, higher genome copies and/or titers of B.1.1.7 and wt-S^B.1.1.7^ compared to wt-S^614G^ were found in the URT (oropharynx, nose) and olfactory bulb. Increased replication and transmission of wt-S ^B.1.1.7^ over wt-S^614G^ were also evident in hACE2-K18Tg mice. Transmission events are rare in mice, however, we observed transmission of B.1.1.7 and wt-S^B.1.1.7^ in 50% of the contact hACE2-K18Tg mice and no detection of wt-S^614G^ in any contact mouse.

### VOC B.1.351 is outcompeted by B.1.1.7 and wt-S^614G^ in vitro and in the hamster model

Neither B.1.1.7 nor B.1.351 demonstrated a replicative fitness advantage compared to wt-S^614G^ in human primary AEC cultures. Furthermore, a clear fitness disadvantage of the B.1.351 variant was particularly apparent in hamsters, where SARS-CoV-2 replicates at high levels in the URT and the lungs. Both wt-S^614G^ and B.1.1.7 outcompeted B.1.351 in terms of replication in experimentally infected hamsters and in transmission to contact animals, where B.1.351 was in the minority by one or two orders of magnitude. This reduced fitness was also evident in previous experiments in K18-hACE2 mice ^18^. The relative reduced intrinsic fitness of B.1.351 in immunologically naïve hosts support the hypothesis that the epidemiological advantage of B.1.351 may be due to immune escape as indicated by reduced efficiency in serum neutralization tests shown here and previously^14^. In immune –convalescent or vaccinated– populations, the immune escape advantage of B.1.351 may prove to be sufficient to compensate for the intrinsic reduced fitness and explains e.g., the low prevalence of this variant in regions with a mainly naïve population.

### B.1.1.7 and wt-S^614G^ equally thrive in the “Superspreader” hamster model

In biochemical assays, we found that the spike protein of B.1.1.7 had a higher affinity than wt-S^614G^ to the hamster ACE2 receptor. Yet, B.1.1.7 and wt-S^614G^ were comparable in their replication and transmission in Syrian hamsters *in vivo* suggesting that, in a model with a high basal rate of replication, the impact of a marginally fitter SARS-CoV-2 variant may not become apparent. Indeed, efficient simultaneous transmission of both variants to contact hamsters was observed in association with high viral loads in infected animals. In models supporting high replication, such as human AEC cultures and hamsters, only major improvement in replication and transmission can be detected when the variants compared already have a very high fitness. In contrast, in ferrets and hACE2-expressing mice, where the bottleneck for viral replication is more stringent, VOCs with a modest, but real increased ability to replicate and transmit can be identified. The similar replication and transmission efficacy in the hamster are also in line with recent publications using the hamster model and VOCs ^19^. The high replication levels observed from the infected hamster, however, might also indicate that the current variants of concern, e.g., B.1.1.7, are on their way to approaching the viral limit for fitness mutations in naïve hosts.

The basal rate of replication is an important factor in the assertion of a variant over a contemporary variant in a naïve population. Some individuals with higher bioaerosol exhalation levels can initiate disproportionate numbers of transmission events, possibly because of higher viral load in the URT, and are therefore called “superspreaders” ^20^. The hamster model might thus resemble the human “superspreader scenario” since there are no clear indications of a predominant variant in this particular competitive transmission experiment. In the ferret and hACE2-KI models, more restrictions are found, e.g., to the URT. Therefore, these models more closely mimic the situation in humans with a dominance of mild infections. While the transmission events were not high overall (3 out of 8 pairs in ferrets, and 4 out of 8 in hACE2-K18Tg mice), the almost exclusive transmission of VOC B.1.1.7 relative to wt-S^614G^ mirrored the higher transmission rate in the human population to some extent, where B.1.1.7 has been responsible for more than 90% of recent infections in most countries in Europe ^21^.

Overall, our study demonstrates that multiple complementary models (hAEC cultures, hamsters, ferrets, homozygous hACE2-KI and hemizygous hACE2-K18Tg mice) are necessary to comprehensively evaluate and fully chisel different aspects of human SARS-CoV-2 infection and the impact of emerging VOCs on the course of the ongoing pandemic. The combined analysis of our results allows a clear conclusion supporting a fitness advantage of B.1.1.7 and a concomitant disadvantage of B.1.351, which is in line with the observed epidemiological dominance and immune escape phenotype of these VOCs. Importantly, and reassuringly, despite the apparent fitness differences of these VOCs, there is no indication of increased pathology nor complete immune escape from humoral immunity.

## Acknowledgments

For their dedicated animal care, we thank Frank Klipp, Doreen Fiedler, Christian Lipinski, and Steffen Kiepert (Friedrich-Loeffler-Institut); K. Sliz, and D. Brechbühl (Institute of Virology and Immunology). For their outstanding technical assistance, we thank Mareen Lange, Christian Korthase, Patrick Zitzow, and Silvia Schuparis (Friedrich-Loeffler-Institut); Gianna Cadau, Paolo Valenti, Barbara Lemaitre (University Hospitals Geneva), Chantal Tougne, Paola Fontannaz and Pascale Sattonnet and Catia Alvarez (University of Geneva). We would like to thank Claire-Anne Siegrist and Arnaud Didierlaurent for providing patient samples; and Angela Huttner and Geraldine Blanchard-Rohner for patient enrollment. We are grateful to S. Berezowska and I. Ramos-Centeno (Institute of Pathology, University of Bern) for providing the tissues via the Tissue Bank Bern. We thank R.G. Crystal (Department of Genetic Medicine, Weill Cornell Medical College, New York NY, USA) for generously providing reagents. We thank M. Schmolke, B. Mazel-Sanchez, and F Abdul (Faculty of Medicine, Geneva) for generously providing A549-hACE2 cells. This work was supported by grants from the Swiss National Science Foundation (SNF), grants no. 31CA30_196062 (CB, RD), 31CA30_196644 (VT, IAE, RD), 310030_173085 (VT), 310030_179260 (RD), 196383 (IAE); the European Commission, Marie Skłodowska-Curie Innovative Training Network ‘HONOURS’, grant agreement no. 721367 (VT, RD); the European Union Project ReCoVer, grant no. GA101003589 (CDr); Core funds of the University of Bern (VT, RD); Core funds of the German Federal Ministry of Food and Agriculture (MB); the Deutsche Forschungsgemeinschaft (DFG), Project no. 453012513 (MB); the Horizon 2020 project “VEO”, grant agreement no. 874735 (MB); COVID-19 special funds from the Swiss Federal Office of Public Health and the Swiss Federal Office of Food Safety and Veterinary Affairs (AS, VT); the Fondation Ancrage Bienfaisance du Groupe Pictet (IAE); the Fondation Privée des Hôpitaux Universitaires de Genève (IAE); the German Ministry of Research, VARIPath, grant no. 01KI2021 (VMC).

## Author contributions

Conceptualization: DH, MB, VT, CB

Data curation: AT, CDe, NJH, LU, JS, RD

Funding acquisition: VT, MB, CB, RD, IE, AS

Investigation: LU, NJH, AT, NE, JS, CDe, BST, BH, MW, MBE, MEL, MLS, DN, VMC, AK, AG, LL, JNK, AB, CW, IBV, MG, KA, BM, CE, LT, MGA, RD, DH

Methodology: BH, AB, FL, JJ

Supervision: CB, VT, MB, DH, RD, IE, CDr

Visualization. AT, CDe, NJH, LU, JS

Writing – original draft: LU, NJH, DH, AT, RD, MB, CB

Writing – review & editing: CB, VT, DH, NH, MB, RD, LT, LU, AB

## Data and materials availability

All data are available in the main text or the supplementary materials.

## Methods

### Cell lines

Vero E6 cells (kindly provided by Doreen Muth, Marcel Müller, and Christian Drosten, Charité, Berlin, Germany) or Vero-TMPRSS2 ^22^ (kindly provided by Stefan Pöhlmann, German Primate Center - Leibniz Institute for Primate Research, Göttingen, Germany) were propagated in Dulbecco’s Modified Eagle Medium-GlutaMAX™ supplemented with 1 mM sodium pyruvate, 10% (v/v) heat-inactivated fetal bovine serum (FBS), 100 μg/ml streptomycin, 100 IU/ml penicillin, 1% (w/v) nonessential amino acids, and 15 mM HEPES (Gibco, Gaithersburg, Maryland, United States of America). Cells were maintained at 37°C in a humidified incubator with 5% CO_2_.

### Viruses

Viruses are listed in Extended Data Table 1 together with the corresponding *in vitro* and *in vivo* experiments in which they were used. Specific amino acid changes are shown schematically in figure S17. Contemporary clinical isolates from the B.1.160 (S^D614G^) (EPI_ISL_414019), B.1.1.7 (EPI_ISL_2131446, EPI_ISL_751799 (L4549)) and B.1.351 (EPI_ISL_803957 (L4550)) were isolated and minimal passaged upon Vero E6 cells. B.1.351 (EPI_ISL_981782) was initially isolated on A549 cells expressing hACE2 before passaging on Vero E6 cells. SARS-CoV-2 B.1.1.7 (L4549) and B.1.351 (L4550) ^18^ were received from the Robert-Koch-Institut Berlin, Germany. Isogenic variants with the B.1.1.7 spike (wt-S^B.1.1.7^) or individual B.1.1.7 spike mutations were introduced into a wild-type SARS-COV-2 “Wuhan” backbone strain comprising the D614G amino acid change (wt-S^614G^), as described previously ^4, 23^. Isogenic viruses were grown on Vero-TMPRSS2 cells, after one passage on human bronchial airway epithelial cells. All viruses were verified by performing whole-genome NGS sequencing (see below). For SARS-CoV-2 B.1.1.7 (L4549), one silent mutation in the ORF1a (sequence position 11741) was determined (C to T with 27% T, 57% strand bias). For SARS-CoV-2 B.1.351 (L4550, available under ENA study accession number PRJEB43810), one nucleotide exchange was detected (A12042C) resulting in the amino acid exchange D3926A in ORF1a and one SNP at sequence position 11730 (C to T with 41%, stand bias 52%).

For all *in vivo* virus competition experiments, we generated inoculum mixtures aiming for a 1:1 ratio of each variant based on virus stock titers. The reported mixture inoculum titers are based on back-titration of the inoculum mixtures and the indicated ratio of each variant was determined by the standard RT-qPCR. SARS-CoV-2 wt-S^614G^ and SARS-CoV-2 B.1.351 (L4550) were used to inoculate hamsters in the wt-S^614G^ vs. B.1.351 study; SARS-CoV-2 B.1.1.7 (L4549), and SARS-CoV-2 B.1.351 (L4550) were used for inoculation in the B.1.1.7 vs. B.1.351 hamster study. SARS-CoV-2 wt-S^614G^, wt-S^B.1.1.7^, B.1.1.7 (L4549), and B.1.351 (L4550) were used to inoculate hACE2 humanized mice in all single virus or mixed virus competition experiments.

### Primary human nasal and bronchial airway epithelial cell (AEC) cultures

Human nasal airway epithelial cell (AEC) cultures were purchased from Epithelix SARL, Geneva, (EP02MP Nasal MucilAir™ Pool of 14 Donors). Bronchial AEC were either established from primary human tracheobronchial epithelial cells isolated from patients (>18 years old) undergoing bronchoscopy or pulmonary resection at the Cantonal Hospital in St. Gallen, Switzerland, or Inselspital in Bern, Switzerland, in accordance with ethical approval (EKSG 11/044, EKSG 11/103, KEK-BE 302/2015, and KEK-BE 1571/2019) or using the basal BCi-NS1.1 cell line with multipotent differentiation capacity (kindly provided by Ronald G. Crystal, Department of Genetic Medicine, Weill Cornell Medical College, New York NY, USA) ^20^. Isolation and maintenance of primary nasal and bronchial AEC cultures were performed according to manufacturer’s guidelines or as previously described ^24, 25^. Individual SARS-CoV-2 infections with contemporary virus isolates were conducted at either 33°C or 37°C as described elsewhere using an MOI of 0.02 ^26^, while all competition experiments and replication kinetics of individual isogenic variants were performed with an MOI of 0.005 as described previously ^24^. The viral load quantification of individual SARS-CoV-2 infections with contemporary virus isolates was performed using the NucliSens easyMAG (BioMérieux) and quantitative real-time PCR (RT-qPCR) targeting the E gene of SARS-CoV-2 as described ^27, 28^. The competition experiments nucleic acids were extracted using the Quick-RNA Viral 96 kit (Zymo research) and the RT-qPCR primers described in Extended Data Table 2. The viral replication of individual isogenic variants was monitored via plaque assay.

### Plaque titration assay

Viruses released into the apical compartments were titrated by plaque assay on Vero E6 cells, as previously described ^24, 29^. Briefly, 2×10^5^ cells/ml were seeded in 24-well plates 1 day prior to titration and inoculated with 10-fold serial dilutions of virus solutions. Inoculums were removed 1 hour post-infection and replaced with overlay medium consisting of DMEM supplemented with 1.2% Avicel (RC-581, FMC biopolymer), 15 mM HEPES, 5 or 10% heat-inactivated FBS, 100 μg/ml streptomycin, and 100 IU/ml penicillin. Cells were incubated at 37°C, 5% CO_2_ for 48 hours, fixed with 4% (v/v) neutral buffered formalin, and stained with crystal violet.

### Serum and plasma samples

Convalescent plasma were collected in the context of a prospective cohort study at the Geneva University Hospitals (HUG) from patients infected with SARS-CoV-2 during the first pandemic wave in 2020, before circulation of any VOC. All patients were confirmed by RT-qPCR for COVID-19; serum or plasma were collected between 30 and 40 days post symptom onsets. Further sera were available through a prospective observational cohort study conducted at Charité-Universitätsmedizin Berlin ^30^, a prospective observational cohort study on immunogenicity and reactogenicity of COVID-19 vaccines in healthcare workers ^31^, and from patients who recovered from mild to moderate COVID-19 for convalescent plasma donation. (https://www.frontiersin.org/articles/10.3389/fimmu.2020.628971/full). Additional analyzes of patient data were performed in accordance with the Berlin State Hospital Law, allowing for pseudonymized scientific analysis of routine patient data.

The studies were approved either by the Geneva Cantonal Ethics Commission (2020-00516) or by the ethics committee of Charité-Universitätsmedizin Berlin (EA2/066/20, EA4/244/20, EA2/092/20, and EA4/245/20).

### Plaque reduction neutralization test (PRNT) (Berlin)

Plaque reduction neutralization tests (PRNT) were done as described before ^32, 33^. Briefly, Vero E6 cells were seeded in a 24-well plate format. Serum or plasma samples were diluted (1:40 to 1:320) and mixed with 100 plaque-forming units of the specific SARS-CoV-2 strain. Each 24-well was incubated with serum/plasma-virus. After 1 hour at 37°C, the supernatants were discarded, and cells were washed once with PBS and supplemented with the overlay solution also used for the plaque assay. After 3 days at 37°C, the supernatants were discarded, and cells were fixed using a 6% formalin solution and stained with crystal violet. All dilutions were tested in duplicates. Serum dilutions with an average plaque reduction of 50% or 90% (PRNT_50/90_) are referred to as titers.

### Plaque reduction neutralization test (PRNT) (Geneva)

Vero-E6 cells were seeded at a density of 4×10^5^ cells/mL in 24-well cell culture plates. All serum/plasma were heat-inactivated at 56°C for 30min and serially diluted in Opti-Pro serum free medium starting from 1:10 until 1:2560 if necessary. Serum/plasma were mixed with 50PFU of variant isolates and incubated at 37°C for 1h. All samples were run in duplicate and for each neutralization experiment an infection control (no serum/plasma) and a reference serum was used to ensure reproducibility between different experiments. Vero-E6 cells were washed 1x with PBS and inoculated with the virus serum/plasma mixture for 1h. Afterwards, the inoculum was removed and 500uL of the overlay medium also used for the plaque assays was added. After incubation for 3 days at 37°C, 5% CO_2_, the overlay medium was removed, cells were fixed in 6% formaldehyde solution for at least 1h, plates were washed 1x with PBS and stained with crystal violet. Plaques were counted in wells inoculated with virus-serum/plasma mixtures and compared to plaque counts in infection control wells. The 90% reduction endpoint titers (PRNT_90_) were calculated by fitting a 4-parameter logistics curve with variable slope to the plaque counts of each serum/plasma.

### Protein expression, purification, and biolayer interferometry assay

SARS-CoV-2 spike protein expression plasmids were constructed to encode the ectodomain of spike protein S^614G^ or S^B.1.1.7^ (residues 1–1208, with a mutated furin cleavage site and K986P/V987P substitutions) followed by a T4 foldon trimerization domain and a polyhistidine purification tag. ACE2 protein (human, hamster, or ferret) expression plasmids were constructed to encode the ectodomain of ACE2 followed by a human IgG1 Fc purification tag. The recombinant proteins were expressed using the Expi293 Expression system (ThermoFisher Scientific) and purified with HisTrap FF columns (for polyhistidine-tagged spike proteins) or with HiTrap Protein A column (for Fc-tagged ACE2 proteins) in FPLC (Cytiva) system.

Binding affinity between the trimeric spike and dimeric ACE2 were evaluated using Octet RED96 instrument at 30°C with a shaking speed at 1000 RPM (ForteBio). Anti-human IgG Fc biosensors (ForteBio) were used. Following 20 minutes of pre-hydration of anti-human IgG Fc biosensors and 1 minute of sensor check, 7.5 nM of human ACE2-Fc, 7.5 nM of hamster ACE2-Fc, or 10 nM of ferret ACE2-Fc in 10X kinetic buffer (ForteBio) were loaded onto the surface of anti-human IgG Fc biosensors for 5 minutes. After 1.5 minutes of baseline equilibration, 5 minutes of association was conducted at 10 – 100 nM S^614G^ or S^B.1.1.7^, followed by 5 minutes of dissociation in the same buffer, which was used for baseline equilibration. For binding assays using ferret ACE2-Fc, association was conducted with 25 – 200 nM S^614G^ or S^B.1.1.7^. The data were corrected by subtracting signal from the reference sample and a 1:1 binding model with global fit was used for determination of affinity constants.

### Animal experimentation ethics declarations

All ferret and hamster experiments were evaluated by the responsible ethics committee of the State Office of Agriculture, Food Safety, and Fishery in Mecklenburg–Western Pomerania (LALLF M-V) and gained governmental approval under registration number LVL MV TSD/7221.3-1-004/21. Mouse studies were conducted in compliance with the Swiss Animal Welfare legislation and approved by the commission for animal experiments of the canton of Bern under license BE-43/20.

### Hamster studies

Six Syrian hamsters (*Mesocricetus auratus*) (Janvier Labs) were inoculated intranasally under a brief inhalation anesthesia with a 70μl mixture of two respective SARS-CoV-2 VOCs (wt-S^614G^ vs. B.1.1.7 mixture, wt-S^614G^ vs. B.1.351 mixture, and B.1.1.7 vs. B.1.351 mixture). Each inoculum was back-titrated and ratios of each variant were determined by RT-qPCR. The wt-S^614G^ and B.1.1.7 mixture held a 1:1.6 ratio with 10^4.3^ TCID_50_ per hamster, the wt-S^614G^ vs. B.1.351 mixture held a 1:3.8 ratio with 10^4.25^ TCID_50_ per hamster, and the B.1.1.7 vs. B.1.351 mixture held a 1.8:1 ratio with 10^5.06^ TCID_50_ per hamster.

Inoculated donor hamsters were isolated in individually ventilated cages for 24 hours. Thereafter, contact hamster I was co-housed to each donor, creating six donor-contact I pairs (Extended Data Fig. 3A). The housing of each hamster pair was strictly separated in individual cage systems to prevent spillover between different pairs. Four days post inoculation (dpi), the individual donor hamsters (inoculated animal) were euthanized. To simulate a second transmission cycle, the original contact hamsters (referred to as contact I) were commingled with a further six naïve hamsters (referred to as contact II), which equates to another six contact I and contact II pairs (Extended Data Fig. 3A). These pairs were co-housed until the end of the study at 21 dpi. Because the first contact hamster (cage 6) in the competition trial wt-S^614G^ vs. B.1.1.7, died already 2 dpc, the second contact hamster for this cage was also co-housed with the donor hamster; thus the first and second contact hamsters in this cage were labeled contact Ia and contact Ib, respectively. To enable sufficient contact between the donor hamster and contact Ib hamster, which was commingled routinely on 4 dpi, the donor hamster was euthanized not before 7 dpi (instead of 4 dpi), when it finally reached the humane end-point criterion for bodyweight (below 80% of 0 dpi body weight).

Viral shedding was monitored by nasal washings in addition to a daily physical examination and body weighing routine. Nasal washing samples were obtained under a short-term isoflurane anesthesia from individual hamsters by administering 200µl PBS to each nostril and collecting the reflux. Animals were sampled daily from 1 dpi to 9 dpi, and afterwards, every other day until 21 dpi. Under euthanasia, serum samples and an organ panel comprising representative upper (URT) and lower respiratory tract (LRT) tissues were collected from each hamster. All animals were observed daily for signs of clinical disease and weight loss. Animals reaching the humane endpoint, e.g., falling below 80% of the initial body weight relative to 0 dpi, were humanely euthanized.

### Ferret studies

Similar to the hamster study, 12 ferrets (six donor ferrets and six transmission I ferrets) from the FLI in-house breeding were housed pairwise in strictly separated cages to prevent spillover contamination. Of these, six ferrets were inoculated with an equal 250 µl mixture of SARS-CoV-2 wt-S^614G^ and B.1.1.7. The inoculum was back-titrated and the ratio of each variant was determined by RT-qPCR. The wt-S^614G^ vs B.1.1.7 mixture held a 1:1.2 ratio with 10^5.875^ TCID_50_ distributed equally into each nostril of donor ferrets. Ferrets were separated for the first 24 hours following inoculation. Subsequently, the ferret pairs were co-housed again, allowing direct contact of donor to contact I ferrets. All ferrets were sampled via nasal washings with 750 µl PBS per nostril under a short-term inhalation anesthesia. Donor ferrets were sampled until euthanasia at 6 dpi, which was followed by the introduction of one additional naïve contact II ferret per cage (n=6), resulting in a 1:1 pairwise setup with contact I and II ferrets (Extended Data Fig. 3B). All ferrets, which were in the study group on the respective days, were sampled on the indicated days. Bodyweight, temperature, and physical condition of all animals were monitored daily throughout the experiment. URT and LRT organ samples, as well as blood samples of all ferrets were taken at respective euthanasia timepoints.

Full autopsy was performed on all animals under BSL3 conditions. The lung, trachea, and nasal conchae were collected and fixed in 10% neutral-buffered formalin for 21 days. The nasal atrium, decalcified nasal turbinates (cross-sections every 3-5 mm), trachea and all lung lobes were trimmed for paraffin embedding. Based on PCR results, tissue sections (3 μm) of all donors (day 6) and one recipient (# 8, day 20) were cut and stained with hematoxylin and eosin (H&E) for light microscopical examination. Immunohistochemistry (IHC) was performed using an anti-SARS nucleocapsid antibody (Novus Biologicals #NB100-56576, dilution 1:200) according to standardized avidin-biotin-peroxidase complex-method producing a red labelling and hematoxylin counterstain. Lung tissue pathology was evaluated according to a detailed score sheet developed by Angele Breithaupt (DipECVP) (SI Table 2). Evaluation and interpretation was performed by board-certified veterinary pathologists (DiplECVP) (AB, IBV).

### Mouse studies

Homozygous hACE2 knock-in mice (B6.Cg-*Ace2^tm1(ACE2)Dwnt^*; henceforth hACE2-KI) and hemizygous transgenic mice (Tg(K18-hACE2)2Prlmn; henceforth hACE2-K18Tg) were described previously ^4, 16^. All mice were produced at the specific-pathogen-free facility of the Institute of Virology and Immunology (Mittelhäusern), where they were maintained in individually ventilated cages (blue line, Tecniplast), with 12-h/12-h light/dark cycle, 22 ± 1 °C ambient temperature and 50 ± 5% humidity, autoclaved food and acidified water. At least 7 days before infection, mice were placed in individually HEPA-filtered cages (IsoCage N, Tecniplast). Mice (10 to 12 weeks old) were anesthetized with isoflurane and infected intranasally with 20 μl per nostril with the virus inoculum described in the results section. One day after inoculation, infected hACE2-K18Tg mice were placed in the cage of another hACE2-K18Tg contact mouse. Mice were monitored daily for bodyweight loss and clinical signs. Oropharyngeal swabs were collected under brief isoflurane anesthesia using ultrafine sterile flock swabs (Hydraflock, Puritan, 25-3318-H). The tips of the swabs were placed in 0.5 ml of RA1 lysis buffer (Macherey-Nagel, ref. 740961) supplemented with 1% β-mercaptoethanol and vortexed. At 2 or 4 dpi, mice were euthanized, and organs were aseptically dissected. Systematic tissue sampling was performed as detailed previously ^4^.

### Animal specimens work up, viral RNA detection, and quantification analyses

Organ samples from ferrets and hamsters were homogenized in a 1 ml mixture composed of equal volumes of Hank’s balanced salts MEM and Earle’s balanced salts MEM containing 2 mM L-glutamine, 850 mg l^−1^ NaHCO3, 120 mg l^−1^ sodium pyruvate, and 1% penicillin–streptomycin) at 300 Hz for 2 min using a Tissuelyser II (Qiagen) and centrifuged to clarify the supernatant. Organ samples from mice were either homogenized in 0.5 mL of RA1 lysis buffer supplemented with 1% β-mercaptoethanol using a Bullet Blender Tissue Homogenizer (Next-Advance Inc.) or in Tube M (Miltenyi Biotech, ref. 130-096-335) containing 1 mL of DMEM using a gentleMACS™ Tissue Dissociator (Miltenyi Biotech). Nucleic acid was extracted from 100 μl of the nasal washes or 200 µL mouse throat swabs after a short centrifugation step or 100 μL of organ sample supernatant using the NucleoMag Vet kit (Macherey Nagel). Nasal washings, throat swabs, and organ samples were tested by RT-qPCR analysis for the ratio of the two different viruses used for inoculation, by applying two different assays, each of them specific for one variant: either the wt-S^614G^, B.1.1.7, or B.1.351 variant (Extended Data Tables 2, 3). Viral RNA copies in swabs and organs in studies using a single variant inoculum in mice were determined using the E protein RT-qPCR exactly as described ^4^.

Four specific RT-qPCR assays for SARS-CoV-2 wt-S^614G^, B.1.1.7, and B.1.351 were designed based on the specific genome deletions within the ORF1 and S gene (Extended Data Table 2). Here, virus specific primers were used to realize a high analytical sensitivity (less than 10 genome copies/µl template) of the according PCR assays, also in samples with a high genome load of the non-matching virus.

The RT-qPCR reaction was prepared using the qScript XLT One-Step RT-qPCR ToughMix (QuantaBio, Beverly, MA, USA) in a volume of 12.5 µl including 1 µl of the respective FAM mix and 2.5 µl of extracted RNA. The reaction was performed for 10 min at 50°C for reverse transcription, 1 min at 95°C for activation, and 42 cycles of 10 sec at 95°C for denaturation, 10 sec at 60°C for annealing and 20 sec at 68°C for elongation. Fluorescence was measured during the annealing phase. All RT-qPCRs were performed on a BioRad real-time CFX96 detection system (Bio-Rad, Hercules, USA). Validation work was performed in comparison to established protocols ^27, 34^.

### Next-generation sequencing

RNA was extracted using the RNAdvance Tissue kit (Beckman Coulter) and the KingFisher Flex System (Thermo Fisher Scientific). Subsequently, RNA was transcribed into cDNA and sequencing libraries were generated as detailed described ^35^ and were sequenced using the Ion Torrent S5XL Instrument (ThermoFisher). Samples with Ct values >20 for SARS-CoV-2 were additionally treated with RNA baits (myBaits, Arbor Biosciences) for SARS-CoV-2 enrichment before sequencing ^36^. Sequence datasets were analyzed via reference mapping with the Genome Sequencer Software Suite (version 2.6; Roche, https://roche.com), default software settings for quality filtering and mapping using EPI_ISL_414019 (B.1), EPI_ISL_2131446 (B.1.1.7) and EPI_ISL_981782 (B.1.351) as references. To identify potential single nucleotide polymorphisms in the read data and for acquisition of variant frequencies for each variant, the variant analysis tool integrated in Geneious Prime (2019.2.3) was applied (default settings, minimum variant frequency 0.02).

### Serological tests of hamsters and ferrets

To evaluate the 100% virus-neutralizing potential of serum samples, we performed a virus neutralization test (VNT) following an established standard protocol as described before, giving the 100% neutralization titers (VNT_100_) ^18^. Briefly, sera were prediluted 1/16 in DMEM and further diluted in log2 steps until a final tested dilution of 1/2048. Each dilution was evaluated for its potential to prevent 10^2^ TCID_50_ SARS-CoV-2 wt-S^614G^, B.1.1.7, and B.1351 from inducing cytopathic effect in Vero E6 cells. Additionally, serum samples from the wt-S^614G^ and B.1.1.7 co-inoculated animals were tested by ELISA for sero-reactivity against the RBD domain ^37^. All samples were generated at the time point of euthanasia of the individual animal.

### Statistical Analysis

Statistical analysis was performed using the GraphPad Prism version 8 or R (version 4.1). Unless noted otherwise, the results are expressed as means ± standard deviation (SD). Two-tailed t-test was used to compare results at different time points post infection, or at two- or four-days post infection; one-way ANOVA with Tukey’s multiple comparisons test was used to compare means of more than 2 groups as indicated in text and figure legends. Significance was defined as p<0.05.

---

**Extended Data Fig. 1.**
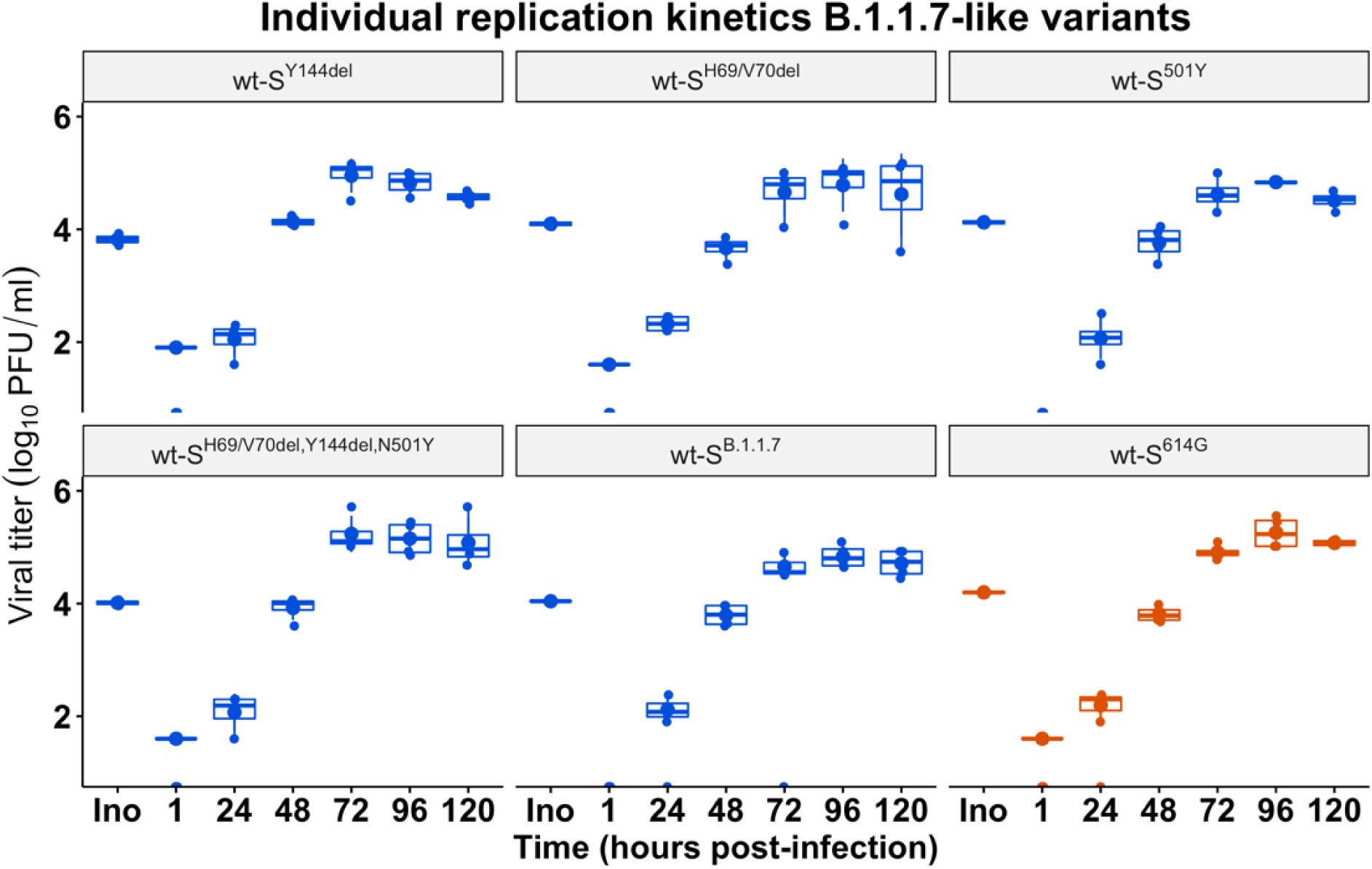
Replication kinetics of B.1.1.7-like SARS-CoV-2. Viral replication kinetics of SARS-CoV-2 B.1.1.7-like variants (blue) and wt-S^614G^ (orange) at 33°C in primary airway epithelial cell (AEC) cultures derived from the bronchial epithelium. Amino acid deletion or substitution in the spike variants are indicated above the corresponding panel. Data are presented as mean values ± standard deviations of four biological replicates, with error bars representing standard deviations.

**Extended Data Fig. 2.**
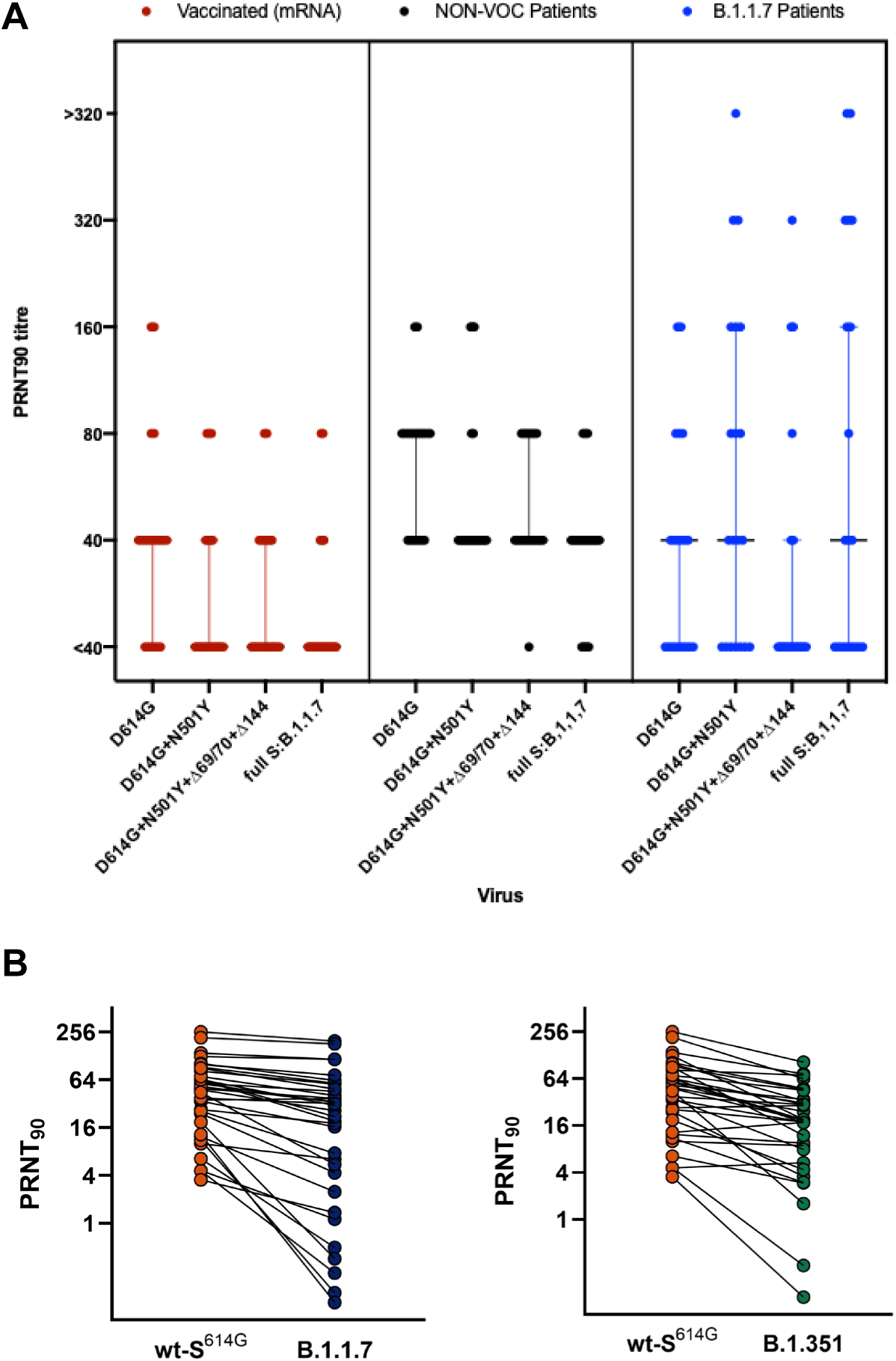
Plaque reduction neutralization tests 90% (PRNT_90_) of patient sera and plasma. (**A**) PRNT_90_ assay of serum from vaccinated, non-VOC patients, and B.1.1.7 patients (N=20/group) for wt-S^614G^ (D614G) and indicated isogenic recombinant mutants encoding some or all indicate B.1.1.7 spike mutations. (**B**) PRNT_90_ assay for convalescent plasma of mild COVID-19 patients (N=34) for clinical isolates wt-S^614G^, B.1.1.7 and B.1.351.

**Extended Data Fig. 3.**
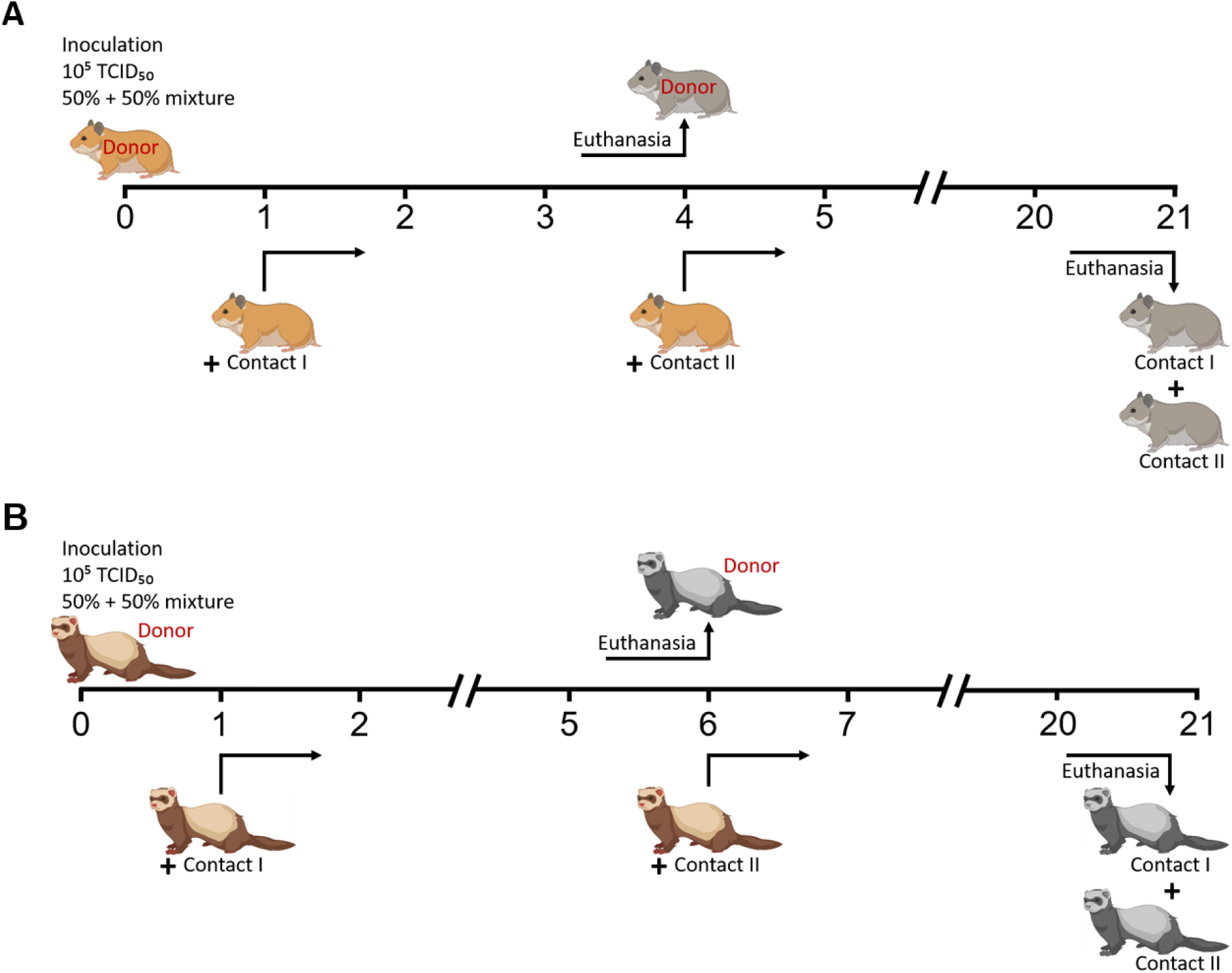
Experimental workflow of competitive transmission experiments in Syrian hamsters and ferrets. (**A**) Timeline of the hamster experiments. Six intranasally inoculated donor hamsters each were co-housed with one naïve contact hamster (1dpi), building six respective donor-contact I pairs. At 4 dpi, the donor hamsters were euthanized and the initial contact hamsters I were co-housed with one additional hamster (Contact II). (**B**) Timeline of the ferret experiment. The scheme was generated with BioRender (https://biorender.com/).

**Extended Data Fig. 4:**
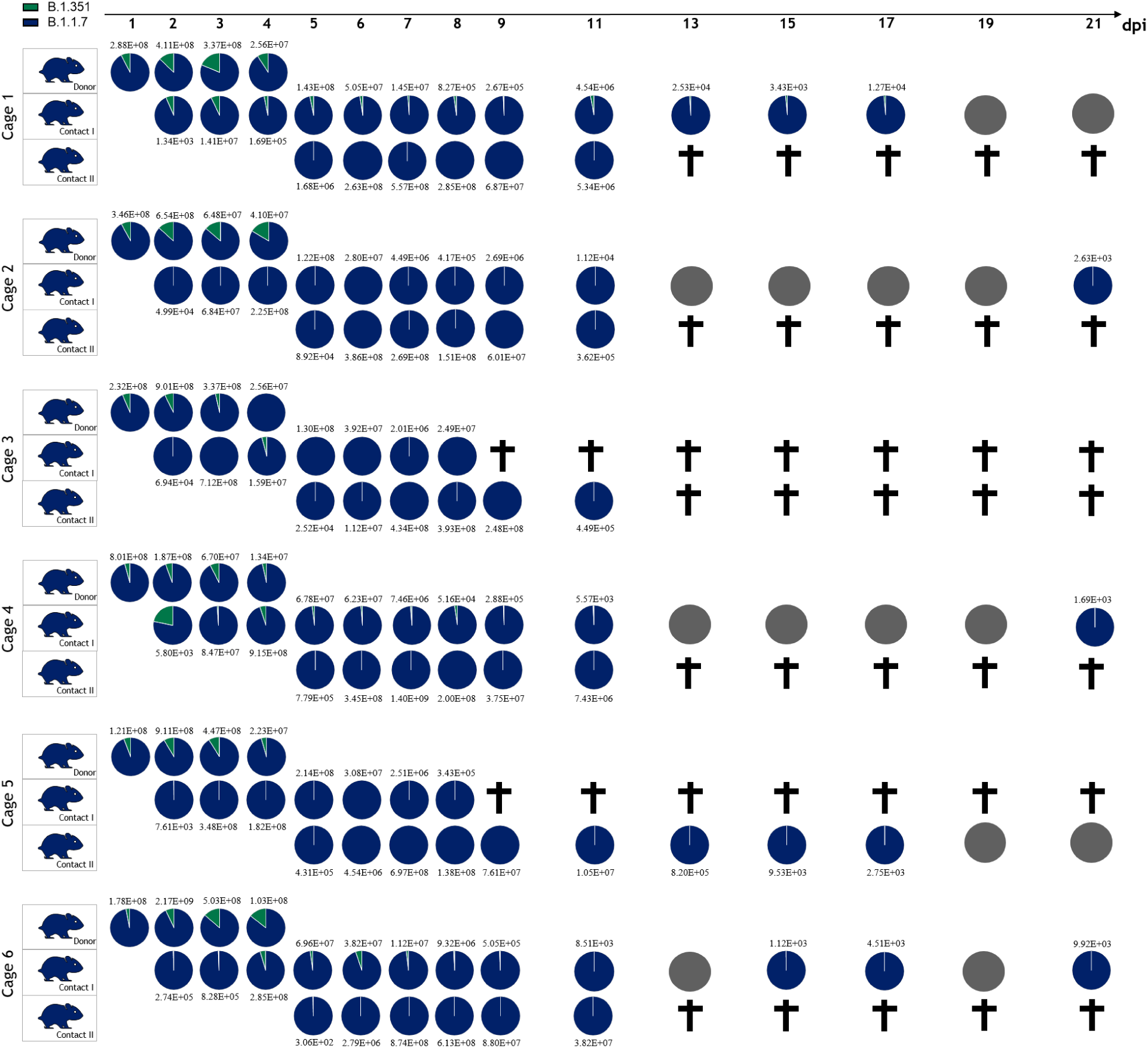
Competitive transmission between B.1.1.7 and B.1.351 in Syrian hamsters. Six donor hamsters were each inoculated with 10^5.06^ TCID_50_ determined by back titration and composed of a mixture of B.1.1.7 (dark blue) and B.1.351 (green) at 1.8:1 ratio determined by RT-qPCR. Donor hamsters, contact I and II hamsters were co-housed sequentially as shown in figure S3A. Nasal washings were performed daily from 1-9 dpi and afterwards every two days until 21 dpi. Each pie chart illustrates the ratio of the respective viruses in nasal washings for each sampling day. Grey pies indicate no detection of viral genomes. Total genome copies/ml are indicated above or below respective pies. Hamster silhouettes are colored according to the dominant variant detected in the latest sample of each animal. Black crosses indicate that the corresponding animal reached the humane endpoint.

**Extended Data Fig. 5.**
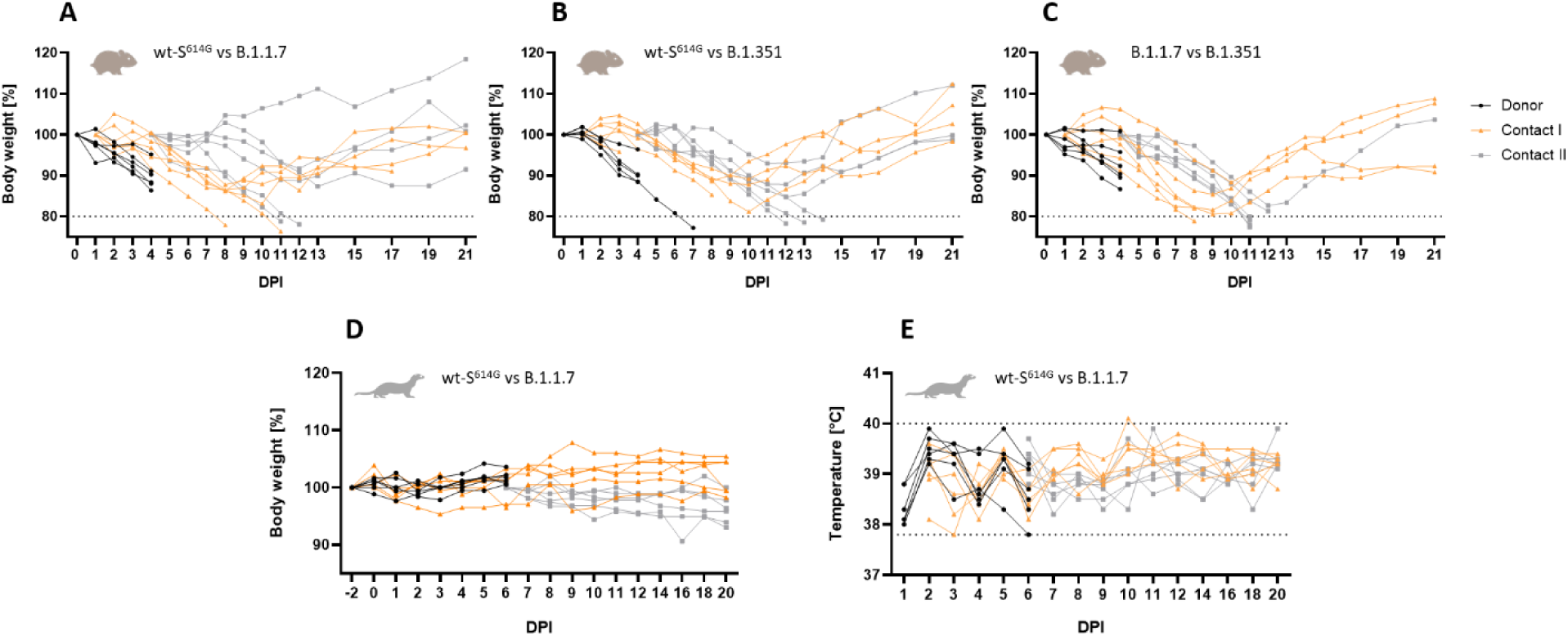
Clinical features of hamsters and ferrets. (**A**-**C**) Syrian hamsters were inoculated with comparable genome equivalent mixture of either wt-S^614G^ and B.1.1.7 (**A**), wt-S^614G^ and B.1.351 (**B**), or B.1.1.7 and B.1.351 (**C)**. In hamsters, body weight was monitored daily until 13 dpi, afterwards every two days until 21 dpi and plotted relative to bodyweight of day 0. The dotted line indicates the humane endpoint criterion of 20% body weight loss from initial bodyweight at which hamsters were promptly euthanized for animal welfare reasons. (**D**, **E**) Ferrets were inoculated intranasally with an equal mixture of wt-S^614G^ and B.1.1.7. Bodyweight (**D**) and temperature (**E**) were monitored daily in ferrets until 12 dpi, and afterwards every 2 days. Grey dotted lines in E indicate the physiologic range for body temperature in ferrets.

**Extended Data Fig. 6.**
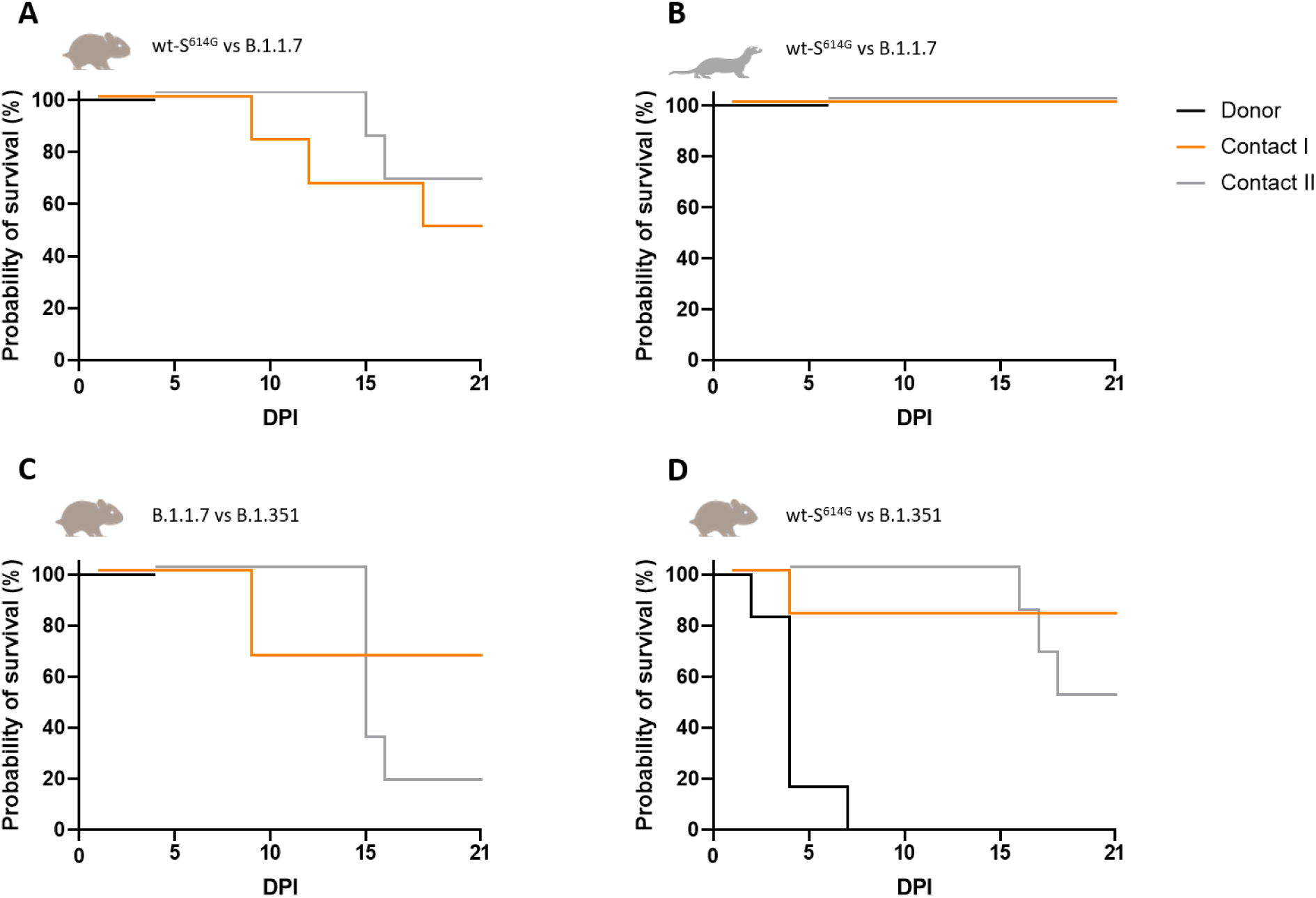
Survival curves for competitive transmission experiments in hamsters and ferrets. Donor hamsters (n=6 per experiment) were euthanized at 4 dpi, donor ferrets (n=6) at 6 dpi. Contact I animals were co-housed at 1 day post inoculation, contact II animals on the day of the respective donor euthanasia; (**A**) Survival of Syrian hamsters during competitive transmission between wt-S^614G^ and B.1.1.7. (**B**) Survival of ferrets during competitive transmission between wt-S^614G^ and B.1.1.7. (**C**) Survival of Syrian hamsters during competitive transmission between B.1.1.7 and B.1.351. (**D**) Survival of Syrian hamsters during competitive transmission between wt-S^614G^ and B.1.351. In the latter study, one donor hamster was kept until it reached the humane endpoint at 7 dpi (instead of the planned euthanasia at 4 dpi) because its co-housed contact I unfortunately passed away during anesthesia at 2 dpi. Additionally, one donor hamster of the same experiment died of the same cause at 2 dpi.

**Extended Data Fig. 7.**
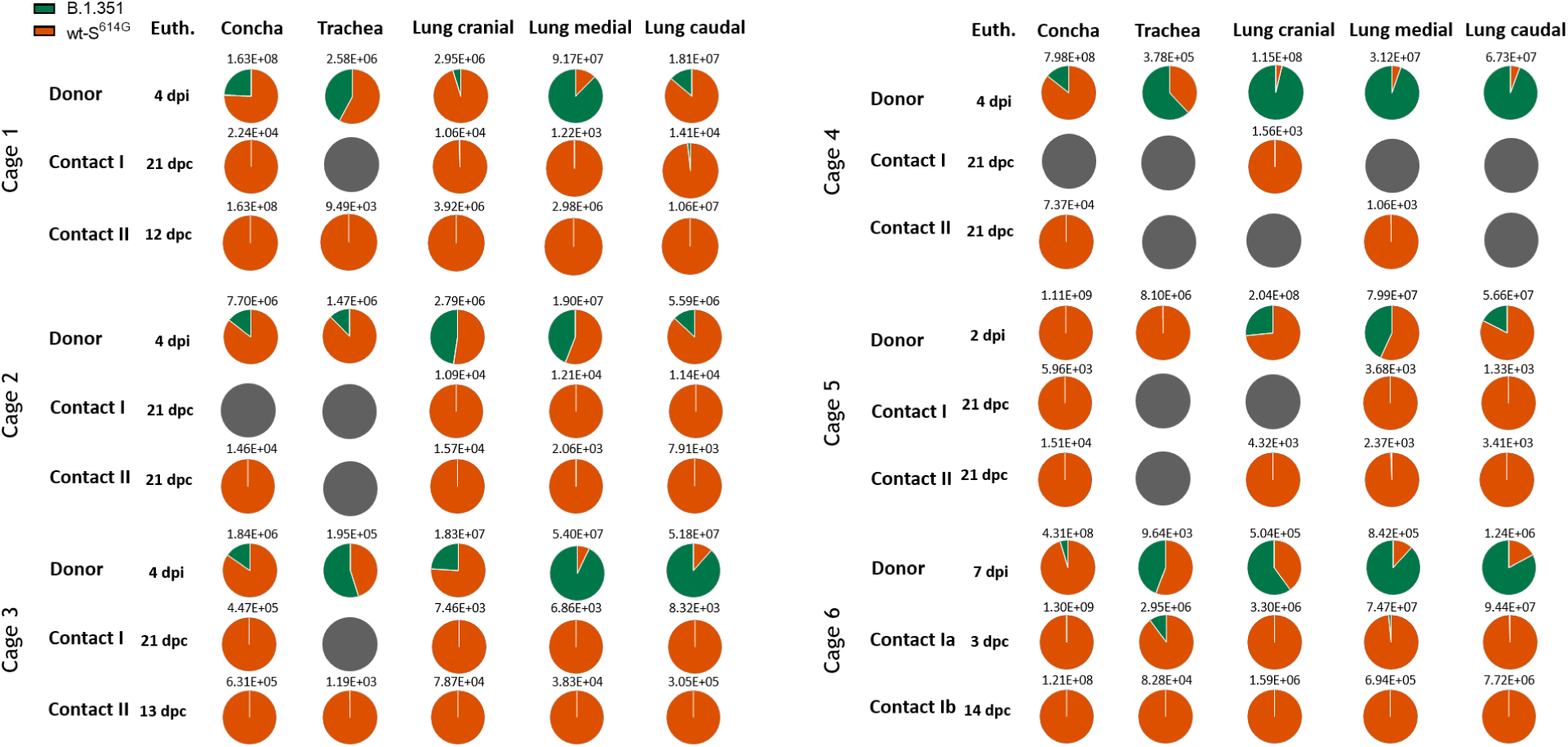
Viral genome load in upper (URT) and lower (LRT) respiratory tract tissues of Syrian hamsters in the competitive transmission experiment between SARS-CoV-2 wt-S^614G^ and B.1.351. Absolute quantification was performed by RT–qPCR analysis of tissue homogenates of donor, contact I and contact II hamsters in relation to a set of defined standards. Tissue samples were collected at the respective time of euthanasia (Euth.). Genome ratios and total amounts of both variants [B.1.351 (green), wt-S^614G^ (orange)] are given for the respective tissues. dpi, days post infection; dpc, days post contact.

**Extended Data Fig. 8.**
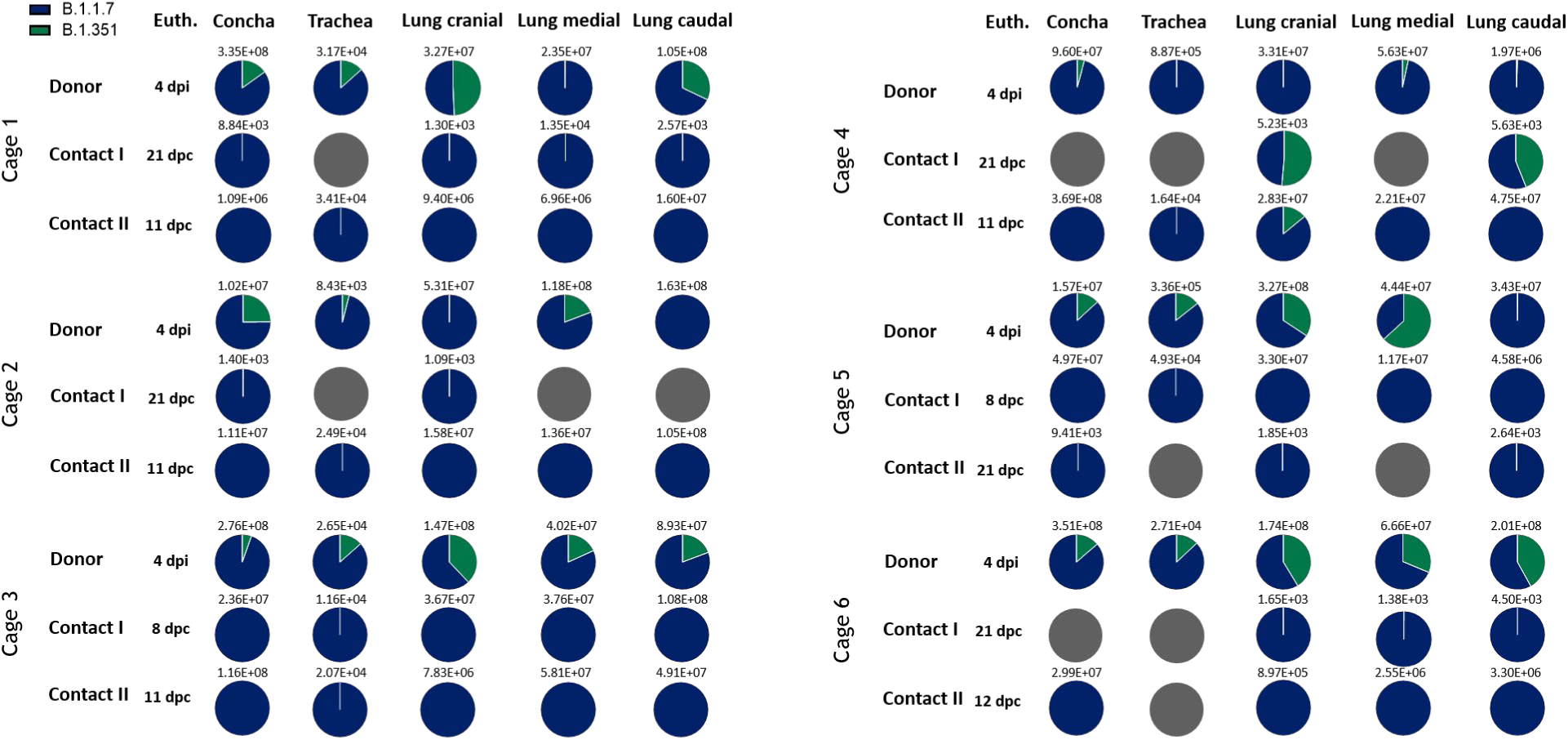
Viral genome load in upper (URT) and lower (LRT) respiratory tract tissues of Syrian hamsters in the competitive transmission experiment between SARS-CoV-2 B.1.1.7 and B.1.351. Absolute quantification was performed by RT–qPCR analysis of tissue homogenates of donor, contact I and contact II hamsters in relation to a set of defined standards. Tissue samples were collected at the respective time of euthanasia (Euth.). Genome ratios and total amounts of both variants [B.1.351 (green), B.1.1.7 (dark blue)] are given for the respective tissues. dpi, days post infection; dpc, days post contact.

**Extended Data Fig. 9.**
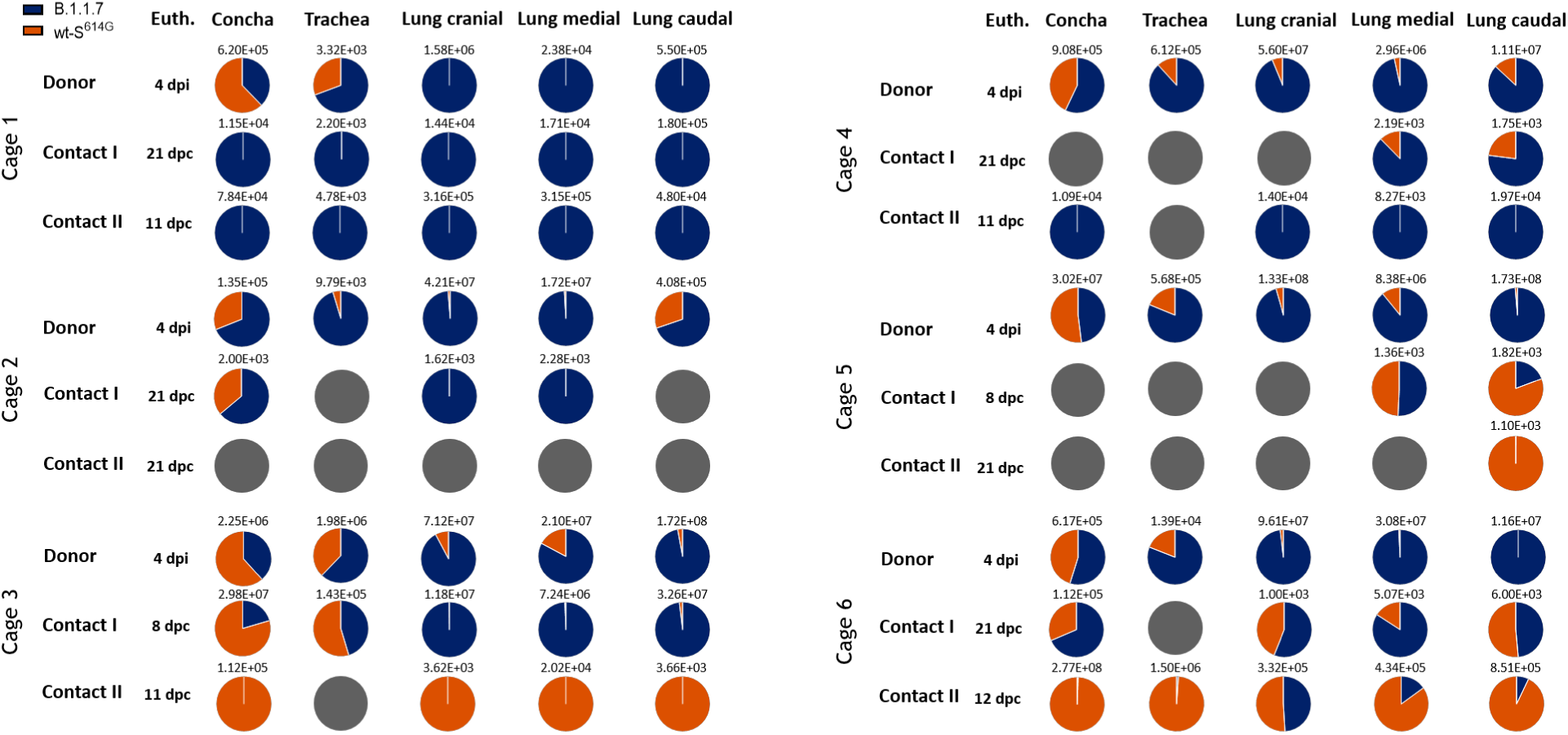
Viral genome load in upper (URT) and lower (LRT) respiratory tract tissues of Syrian hamsters in the competitive transmission experiment between SARS-CoV-2 B.1.1.7 and wt-S^614G^. Absolute quantification was performed by RT–qPCR analysis of tissue homogenates of donor, contact I and contact II hamsters in relation to a set of defined standards. Tissue samples were collected at the respective time of euthanasia (Euth.). Genome ratios and total amounts of both variants [B.1.1.7 (dark blue), wt-S^614G^ (orange)] are given for the respective tissues. dpi, days post infection; dpc, days post contact.

**Extended Data Fig. 10.**
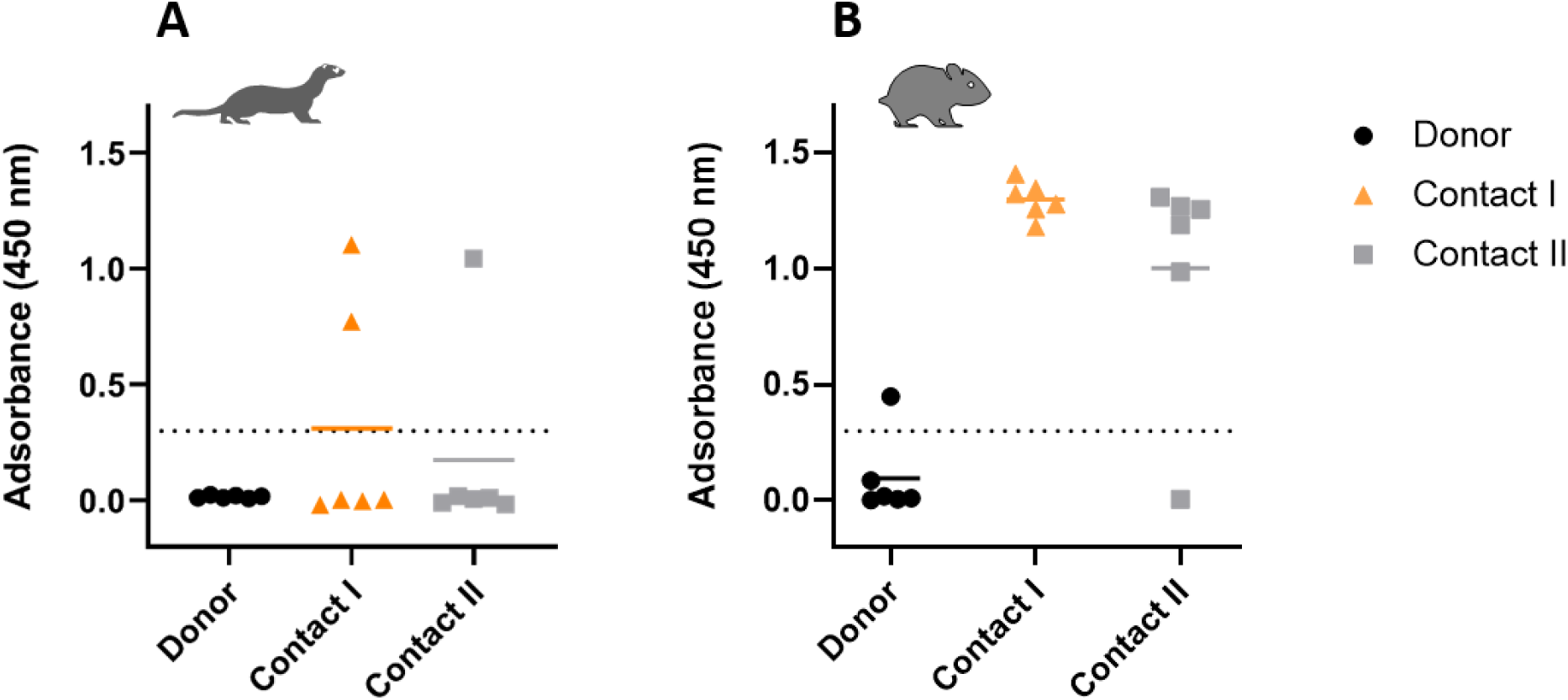
Indirect ELISA against the RBD of SARS-CoV-2. Sera of ferrets (**A**) and hamsters (**B**) inoculated with a mixture of wt-S^614G^ and B.1.1.7 were tested for specific reactivity against the SARS-CoV-2 RBD-SD1 domain (wt-S amino acids 319-519) at the terminal endpoints for each animal.

**Extended Data Fig. 11.**
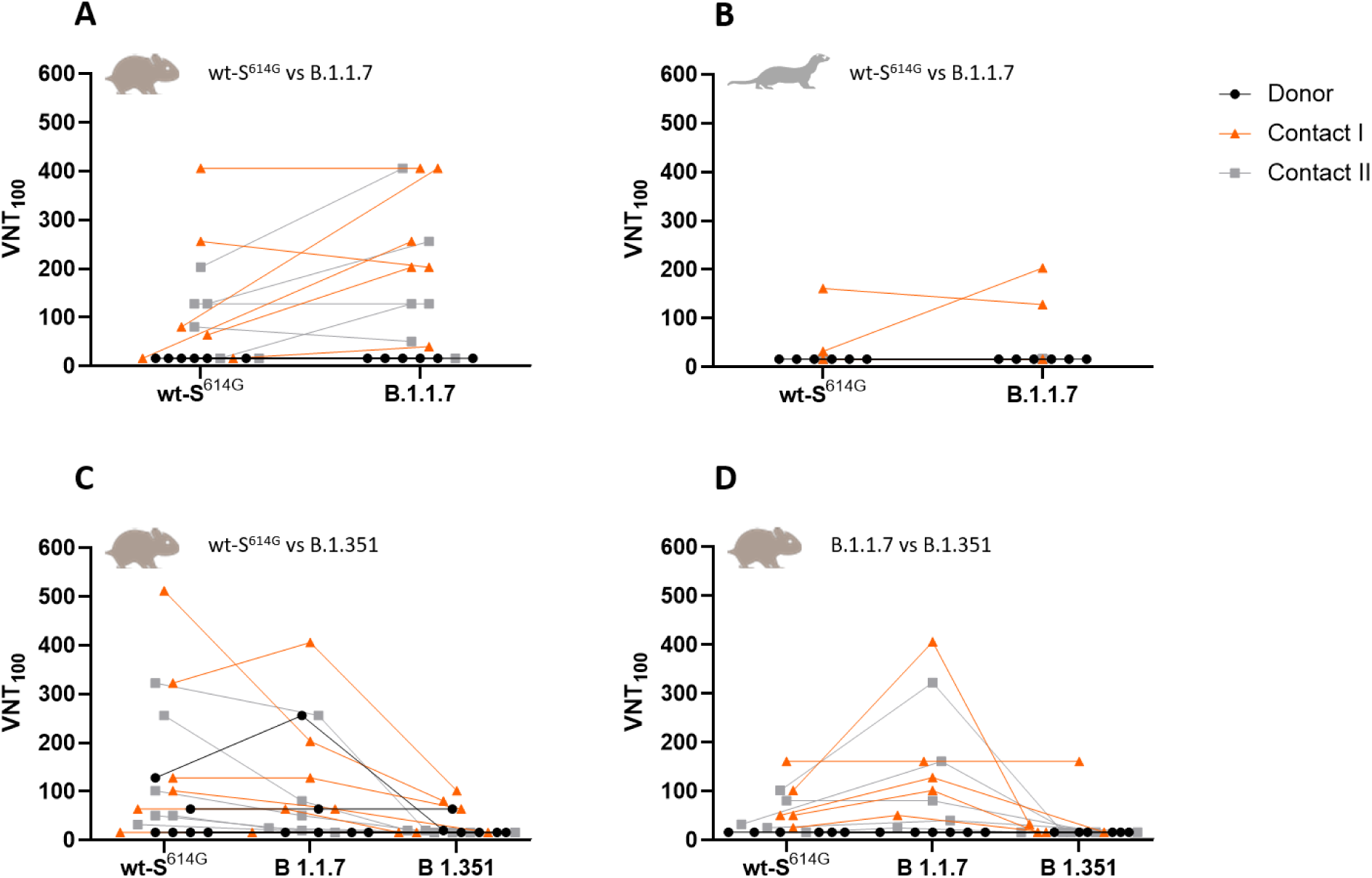
Virus neutralization titers of animal sera. SARS-CoV-2 100% neutralization titers (VNT_100_) in serum samples (log2 titer) of sera from wt-S^614G^ and B.1.1.7, co-inoculated hamsters (**A**) and ferrets (**B**), wt-S^614G^ and B.1.351 inoculated hamsters (**C**) and B.1.1.7 and B.1.351 inoculated hamsters (**D**). Sera from all donor, contact I and contact II hamsters (**A**) and ferrets (**B**) used in the competitive transmission experiment between wt-S^614G^ and B.1.1.7 was collected at respective timepoints of euthanasia and were tested against SARS-CoV-2 wt-S^614G^ and B.1.1.7 on VeroE6 cells. The sera of the donor, contact I and contact II hamsters used in the competitive transmission experiment between B.1.351 and wt-S^614G^ (**C**), as well as the sera of the B.1.351 and B.1.1.7 co-inoculated hamsters (**D**) were tested against wt-S^614G^, B.1.1.7 and B.1.351.

**Extended Data Fig. 12.**
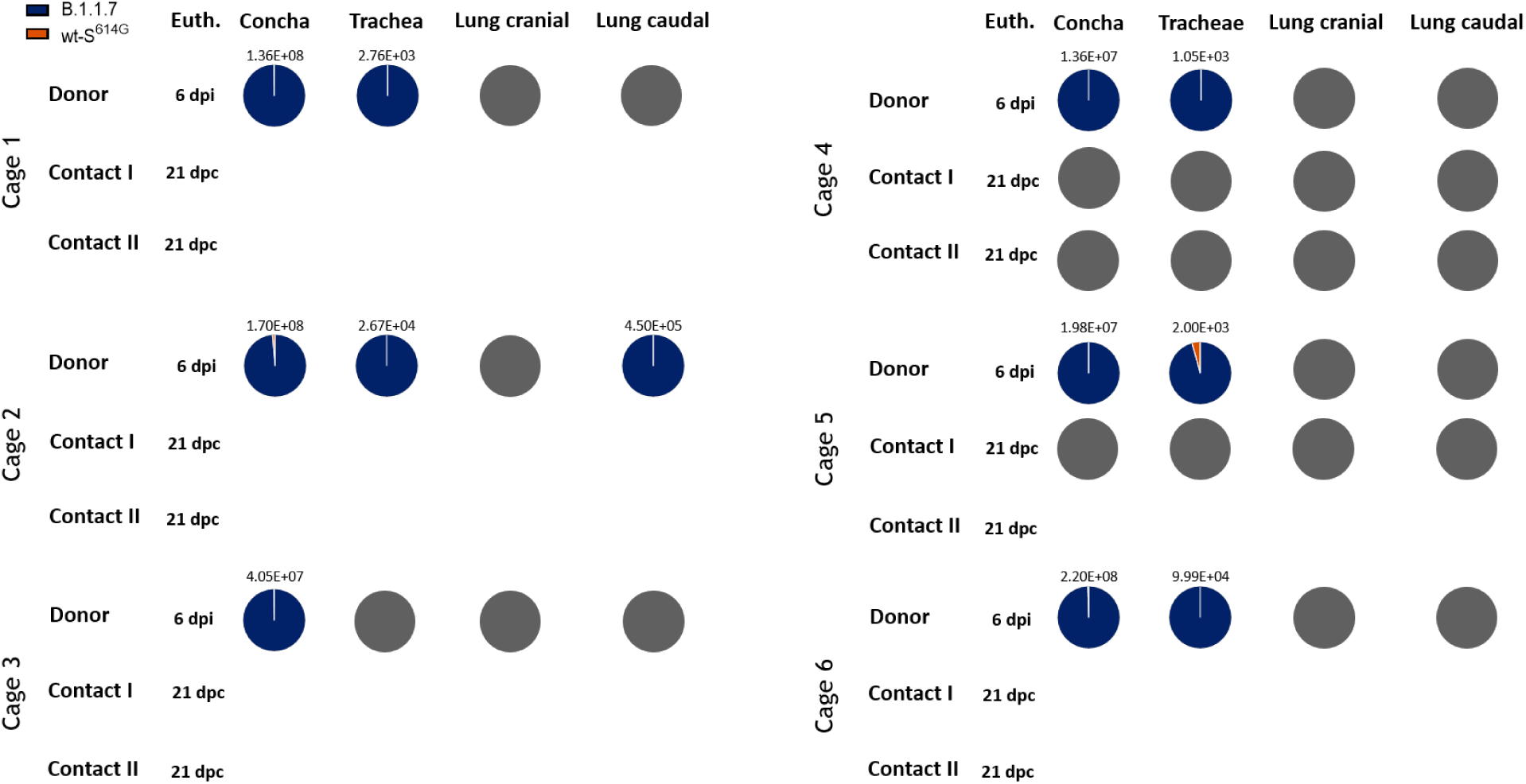
Viral genome load in upper (URT) and lower (LRT) respiratory tract tissue of ferrets in the competitive transmission experiment between SARS-CoV-2 B.1.1.7 and wt-S^614G^. Absolute quantification was performed by RT–qPCR analysis of tissue homogenates of donor, contact I and contact II ferrets in relation to a set of defined standards. Tissue samples were collected at the respective time of euthanasia (Euth.). Genome ratios and total amounts of both variants [B.1.1.7 (blue), wt-S^614G^ (orange)] are given for the respective tissues. dpi, days post infection; dpc, days post contact.

**Extended Data Fig. 13.**
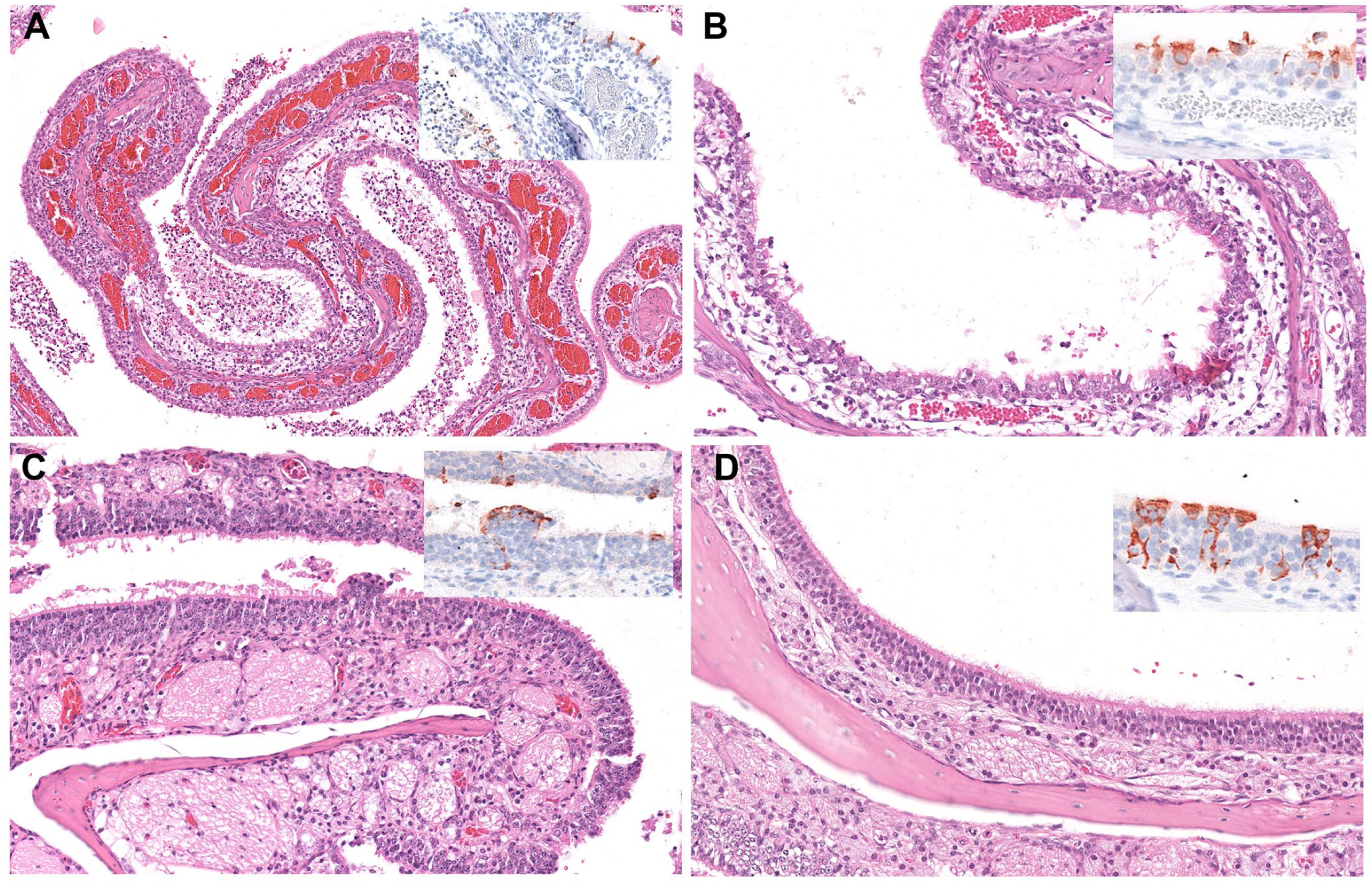
Histopathology of nasal conchae of donor ferrets inoculated with wt-S^614G^ and B.1.1.7. Representative micrographs of hematoxylin and eosin staining of 3 μm sections of nasal conchae of donor ferrets 6 dpi. Insets show immunohistochemistry staining of SARS-CoV-2 with anti-SARS nucleocapsid antibody with hematoxylin counterstain. **(A-D)** The respiratory (A, B) and olfactory (C, D) nasal mucosa exhibited rhinitis with varying severity. Lesion-associated antigen was found in ciliated cells of the respiratory epithelium (A, B) and in sustentacular cells of the olfactory epithelium (C, D) in all animals at 6 dpi.

**Extended Data Fig. 14.**
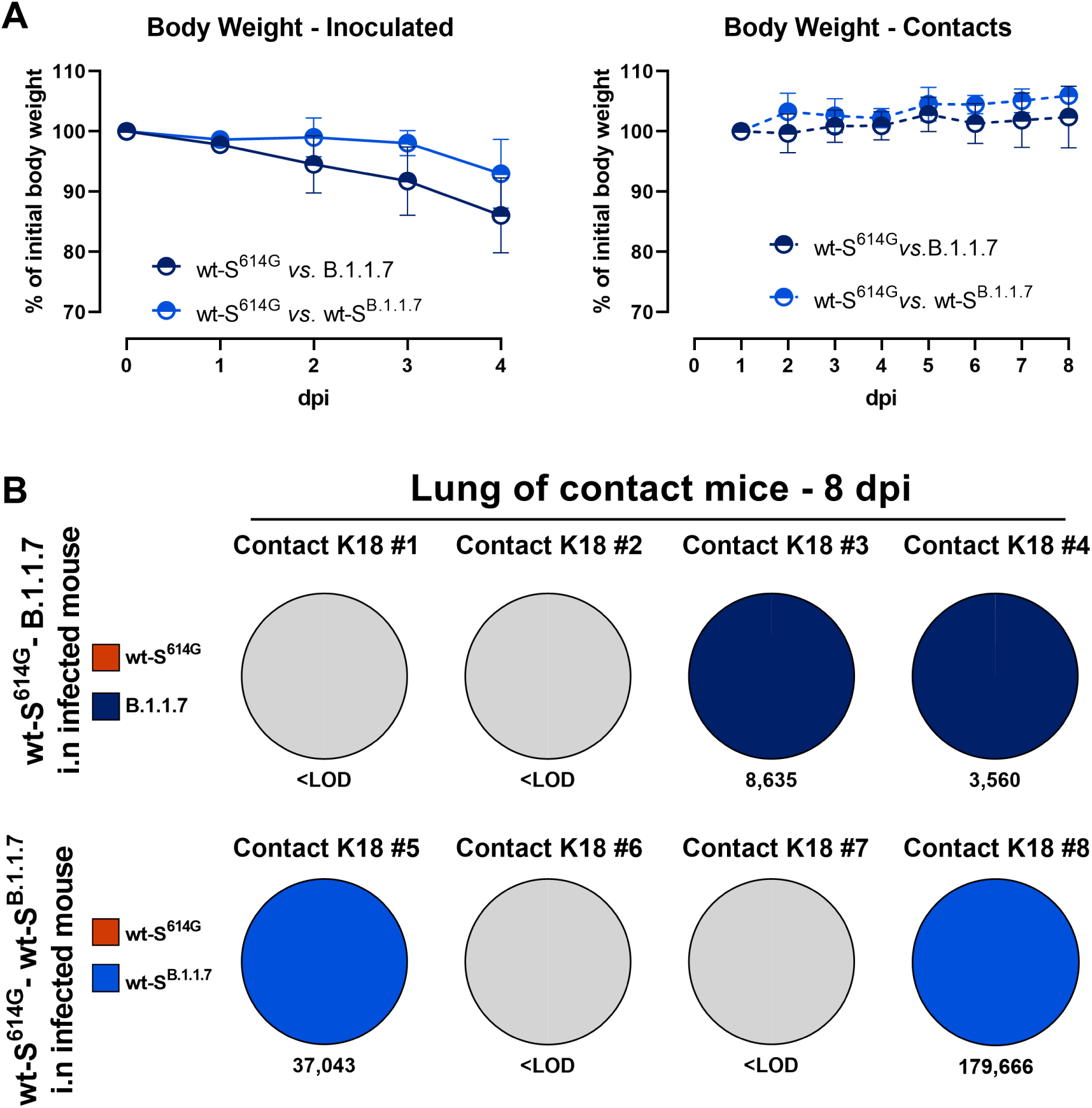
Bodyweight and transmission in hACE2-K18Tg mice. hACE2-K18Tg mice inoculated with a mixture of wt-S^614G^ and B.1.1.7, or wt-S^614G^ and wt-S^B.1.1.7^. (**A**) Relative body weight in donor mice (left panel), and in contact mice (right panel) (n=4/group). (**B**) Pie chart illustrating the ratio of wt-S^614G^ (orange) with B.1.1.7 (dark blue), or with wt-S^B.1.1.7^ (light blue) in corresponding experiments in lung homogenates of contact mice at 7 dpc. Samples below 10^3^ viral genome copies per organ were considered under the limit of detection (<LOD).

**Extended Data Fig. 15.**
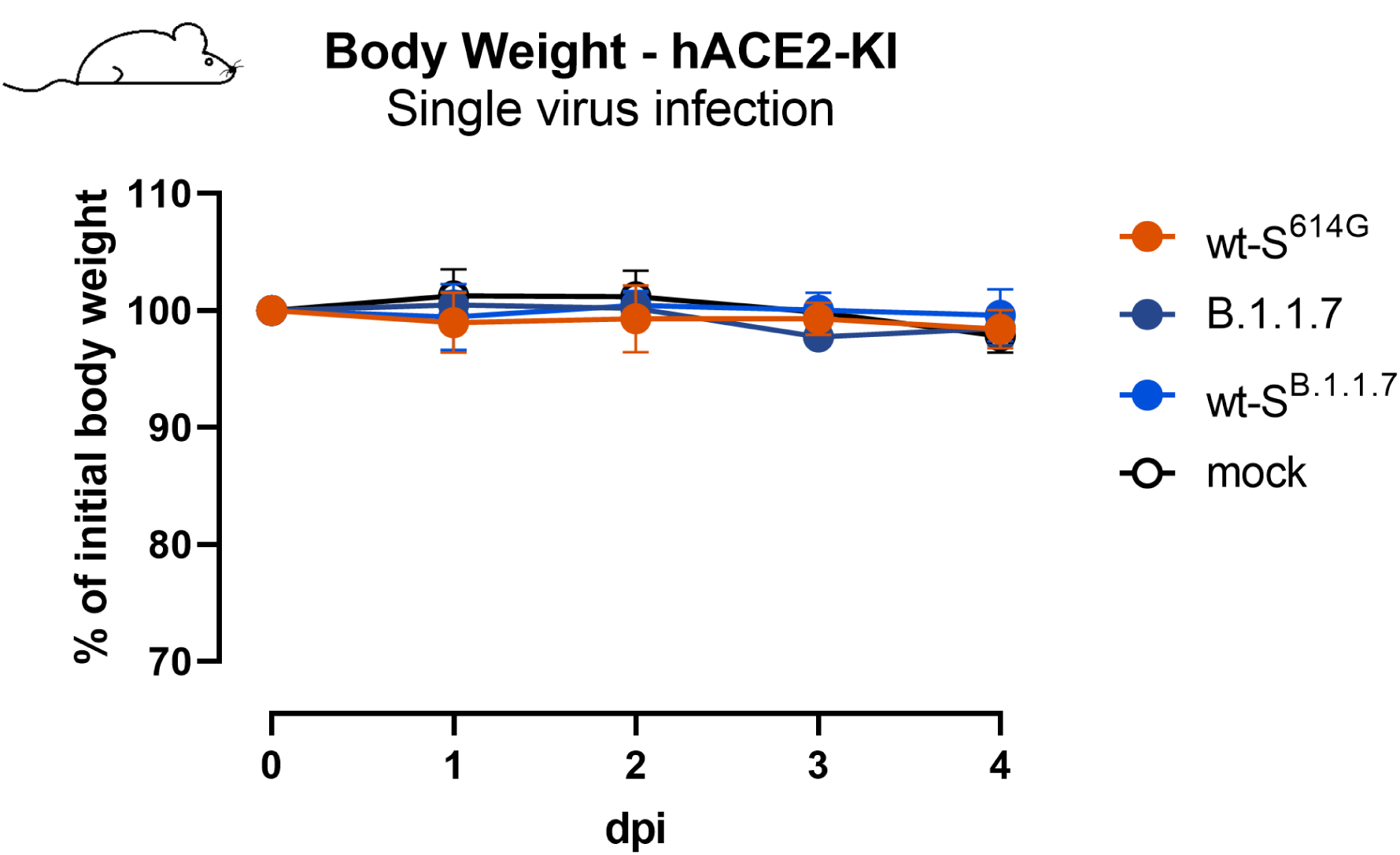
Bodyweight of hACE2-KI mice. Mice mock-infected or inoculated with 10^4^ PFU of wt-S^614G^, B.1.1.7, or wt-S^B.1.1.7^ (n = 8 until 2 dpi, and n= 4 from 3 dpi).

**Extended Data Fig. 16.**
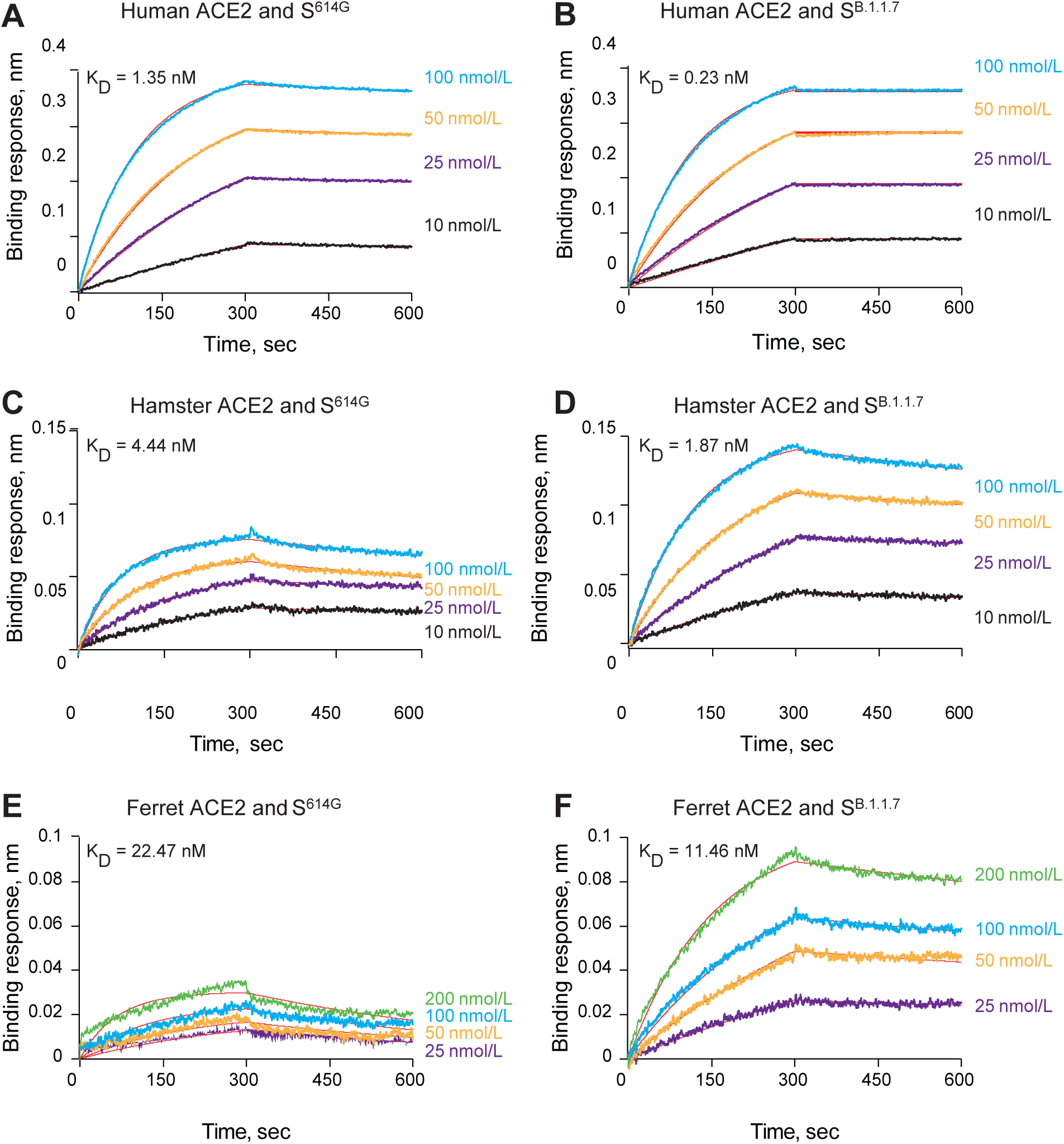
Binding of spike variants with ACE2 of different species. Affinity between spike protein (S^614G^ or S^B.1.1.7^) and (**A, B**) human, (**C, D**) hamster, and ferret (**E, F**) ACE2 determined by Bio-layer interferometry. Human, hamster, or ferret ACE2 with IgG1 Fc tag were loaded on Anti-human IgG Fc biosensors and binding kinetics were conducted using different concentrations of spike trimer proteins (10 nM -100 nM for human or hamster ACE2 and 25 nM to 200 nM for ferret ACE2).

**Extended Data Fig. 17.**
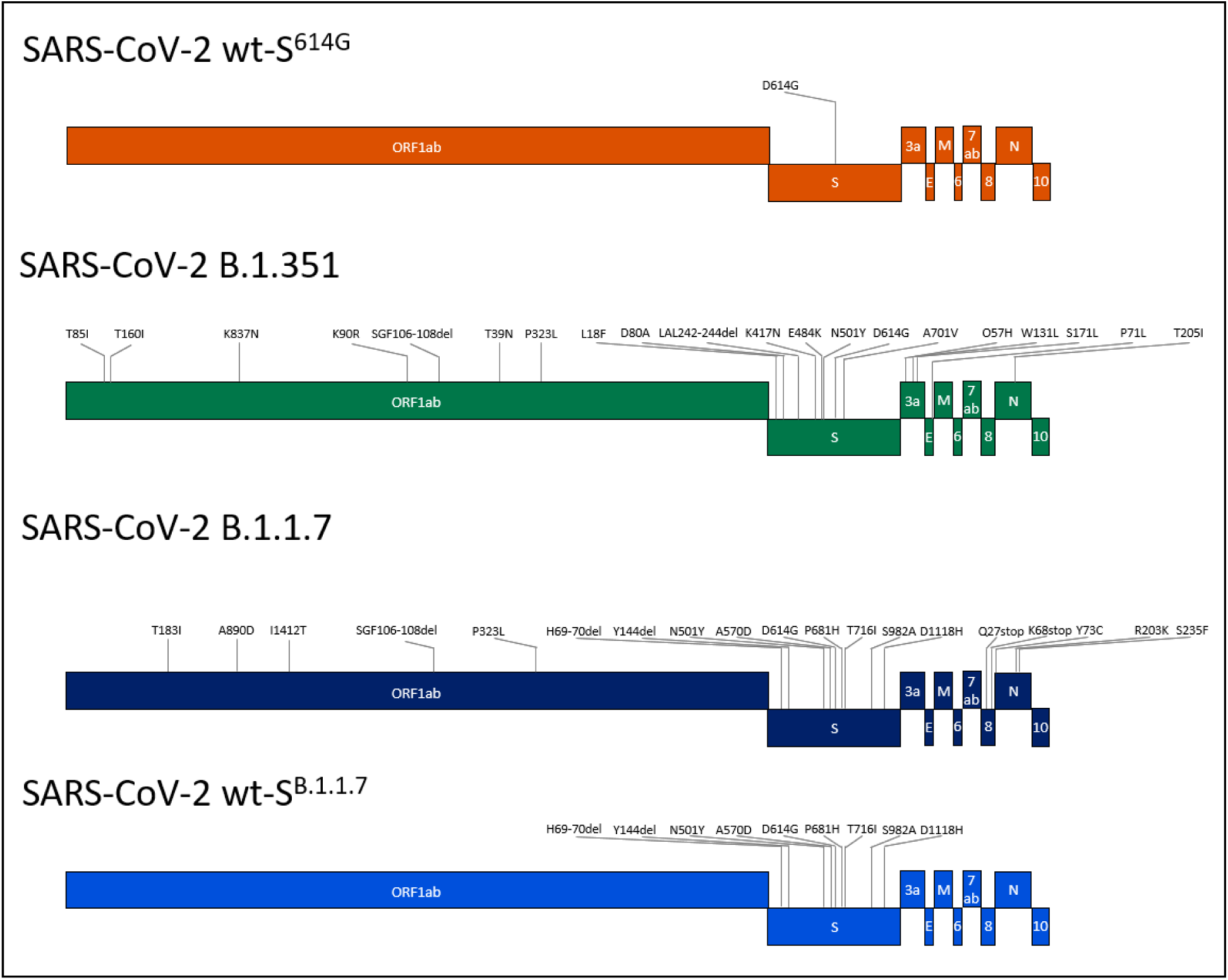
Genome sequences of used SARS-CoV-2 variants. Colors of the variants represent respective viruses in the different experiments. Grey lines indicate positions of known mutations of each virus strain.

**Extended Data Table 1.** Sequence mutations in S in SARS-CoV-2 recombinant strains. (*) SARS-CoV-2 wt-S^614G^ used for the competitive experiments between wt-S^614G^ vs. B.1.1.7 in hamsters and ferrets had an exchange in 54% of the analyzed contigs from C to A in position 23615, located in the S gene.

| SARS-CoV-2 viruses | Mutations in S | Use | Accession-no. / Study-no. /GISAID |
| --- | --- | --- | --- |
| **<u>Isolates:</u>** |  |  |  |
| wt-S<sup>614G</sup><br>(B.1.610) | D614G | Kinetics in vitro | EPI_ISL_414019 |
| B.1.1.7 | H69/V70del, Y144del, N501Y, A570D, D614G, P681H, T716I, S982A, D1118H | Kinetics in vitro | EPI_ISL_2131446 |
| B.1.351 | L18F, D80A, D215G, L242/A243/L244del, K417N, E484K, N501Y, D614G, A701V, | Kinetics in vitro | EPI_ISL_981782 |
| B.1.1.7 (L4549) | H69/V70del, Y144del, N501Y, A570D, D614G, P681H, T716I, S982A, D1118H | competition wt-S<sup>614G</sup> vs B.1.1.7; hamster, ferret, and mice<br>Single virus infection, mice | PRJEB45736, L4549<br>EPI_ISL_803957 |
| B.1.351 (L4550) | L18F, D80A, D215G, L242/L243/A244del, K417N, E484K, N501Y, D614G, A701V | competition wt-S<sup>614G</sup> vs B.1.351 and B.1.1.7 vs B.1.351; hamster <sup>18</sup> | PRJEB43810, L4550, EPI_ISL_751799, |
| **<u>Recombinant clones:</u>** |  |  |  |
| wt-S<sup>614G</sup> | D614G | competition wt-S<sup>614G</sup> vs B.1.351; hamster<br>competition wt-S<sup>614G</sup> vs B.1.1.7; mice<br>competition wt-S<sup>614G</sup> vs wt-S<sup>B.1.1.7</sup>; mice<br>Single virus infection, mice<br>Competition with wt-S<sup>614G</sup> in vitro | MT108784 <sup>4</sup> |
| wt-S<sup>614G</sup> (*)<br>(L4595) | D614G, R685S | competition wt-S<sup>614G</sup> vs B.1.1.7; hamster and ferret | PRJEB45736, L4595 |
| wt-S<sup>B.1.1.7</sup> | H69/V70del, Y144del, N501Y, A570D, D614G, P681H, T716I, S982A, D1118H | competition wt-S<sup>614G</sup> vs wt-S<sup>B.1.1.7</sup>; mice<br>Single virus infection, mice |  |
| wt-S<sup>H69/V70del</sup> | H69/V70del, D614G | Competition with wt-S<sup>614G</sup> in vitro |  |
| wt-S<sup>Y144del</sup> | Y144del, D614G | Competition with wt-S<sup>614G</sup> in vitro |  |
| wt-S<sup>N501Y</sup> | N501Y, D614G | Competition with wt-S<sup>614G</sup> in vitro |  |
| wt-S<sup>H69/V70del,Y144del,N501Y</sup> | H69/V70del, Y144del, N501Y, D614G | Competition with wt-S<sup>614G</sup> in vitro |  |

**Extended Data Table 2.**
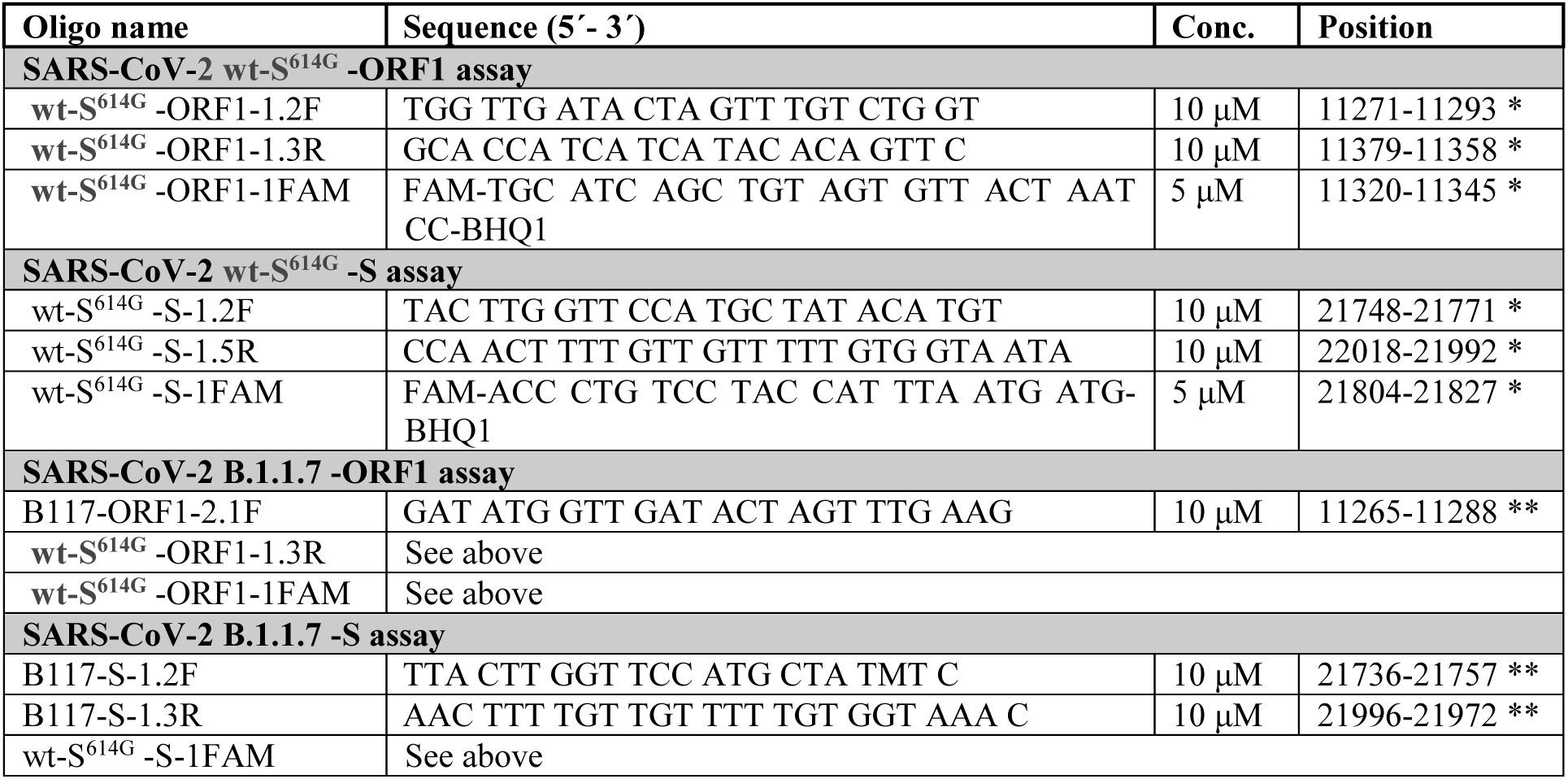
Sequences of primer and probes for RT-qPCR assays. * Position based on NC_045512; ** Position based on MW963651. Conc, concentration.

**Extended Data Table 3.** Attribution of RTqPCR assays used for the individual competitive transmission experiments.

| Assay | For detection of |
| --- | --- |
| **wt-S<sup>614G</sup> vs B.1.1.7 (Hamsters and ferrets)** |  |
| SARS-CoV-2 wt-S<sup>614G</sup> -ORF1 assay | SARS-CoV-2 wt-S<sup>614G</sup> |
| SARS-CoV-2 B.1.1.7 -ORF1 assay | SARS-CoV-2 B.1.1.7 |
| **wt-S<sup>614G</sup> vs B.1.1.7 (Mouse studies and human AEC cultures)** |  |
| SARS-CoV-2 wt-S<sup>614G</sup> -S assay | SARS-CoV-2 wt-S<sup>614G</sup> |
| SARS-CoV-2 B.1.1.7 -S assay | SARS-CoV-2 B.1.1.7 |
| SARS-CoV-2 B.1.1.7 -S assay | SARS-CoV-2 wt-S<sup>B.1.1.7</sup> |
| **wt-S<sup>614G</sup> vs B.1.351 (Hamsters)** |  |
| SARS-CoV-2 wt-S<sup>614G</sup> -ORF1 assay | SARS-CoV-2 wt-S<sup>614G</sup> |
| SARS-CoV-2 B.1.1.7 -ORF1 assay | SARS-CoV-2 B.1.351 |
| **B.1.1.7 vs B.1.351 (Hamsters)** |  |
| SARS-CoV-2 B.1.1.7 -S assay | SARS-CoV-2 B.1.1.7 |
| SARS-CoV-2 wt-S<sup>614G</sup> -S assay | SARS-CoV-2 B.1.351 |

## Legends for Supplementary Information Tables

Supplementary Information Table 1. Histopathological score of lungs of hACE2-KI mice infected with single virus inoculum of wt-S^614G^, B.1.1.7, or wt-S^B.1.1.7^. Hematoxilin and eosin stained slides of left lungs of hACE2-KI mice were scored by a board-certified veterinary pathologist, who was blinded to the identity of the specimen. The scoring scheme is detailed in SI Table 2.

Supplementary Information Table 2. Lung pathology scoring scheme.

