## Supplementary Information Table 1 for "Enhanced fitness of SARS-CoV-2 variant of concern B.1.1.7, but not B.1.351, in animal models"

| | Mock | | | | | | wt-$S^{614G}$ | | | | | | | | B.1.1.7 | | | | | | | | wt-$S^{B.1.1.7}$ | | | | | | | |
|---|---|---|---|---|---|---|---|---|---|---|---|---|---|---|---|---|---|---|---|---|---|---|---|---|---|---|---|---|---|---|
| Days post-inoculation | 2 | 2 | 2 | 4 | 4 | 4 | 2 | 2 | 2 | 2 | 4 | 4 | 4 | 4 | 2 | 2 | 2 | 2 | 4 | 4 | 4 | 4 | 2 | 2 | 2 | 2 | 4 | 4 | 4 | 4 |
| Mouse | M1 | M2 | M3 | M4 | M5 | M6 | KI #1 | KI #2 | KI #3 | KI #4 | KI #5 | KI #6 | KI #7 | KI #8 | KI #9 | KI #10 | KI #11 | KI #12 | KI #13 | KI #14 | KI #15 | KI #16 | KI #17 | KI #18 | KI #19 | KI #20 | KI #21 | KI #22 | KI #23 | KI #24 |
| **Overview** | | | | | | | | | | | | | | | | | | | | | | | | | | | | | | |
| Alveolar edema | 0 | 0 | 0 | 0 | 0 | 0 | 0 | 0 | 0 | 0 | 0 | | 0 | 0 | 0 | 0 | 0 | 0 | 0 | 0 | 0 | 0 | 0 | 0 | 0 | 1 | 0 | 0 | 0 | 0 |
| Emphysema | 0 | 0 | 0 | 0 | 0 | 0 | 0 | 0 | 0 | 0 | 0 | | 0 | 0 | 0 | 0 | 0 | 0 | 0 | 0 | 0 | 0 | 0 | 0 | 0 | 0 | 0 | 0 | 0 | 0 |
| Atelectasis | 0 | 0 | 0 | 0 | 0 | 0 | 0 | 0 | 0 | 0 | 0 | | 0 | 0 | 0 | 0 | 0 | 0 | 0 | 0 | 0 | 0 | 0 | 0 | 0 | 0 | 0 | 0 | 0 | 0 |
| **Higher magnification** | | | | | | | | | | | | | | | | | | | | | | | | | | | | | | |
| **Inflammation score** | | | | | | | | | | | | | | | | | | | | | | | | | | | | | | |
| Alveolar infiltrates | 0 | 0 | 0 | 0 | 0 | 0 | 0 | 0 | 0 | 0 | 0 | | 0 | 0 | 0 | 0 | 0 | 0 | 0 | 0 | 0 | 0 | 0 | 0 | 0 | 1 | 0 | 0 | 0 | 0 |
| Grade | 0 | 0 | 0 | 0 | 0 | 0 | 0 | 0 | 0 | 0 | 0 | | 0 | 0 | 0 | 0 | 0 | 0 | 0 | 0 | 0 | 0 | 0 | 0 | 0 | 1 | 0 | 0 | 0 | 0 |
| Predominant inflammatory cell type | 0 | 0 | 0 | 0 | 0 | 0 | 0 | 0 | 0 | 0 | 0 | | 0 | 0 | 0 | 0 | 0 | 0 | 0 | 0 | 0 | 0 | 0 | 0 | 0 | M | 0 | 0 | 0 | 0 |
| Interstitial infiltrates | 0 | 0 | 0 | 0 | 2 | 0 | 0 | 0 | 0 | 0 | 0 | | 0 | 1 | 0 | 0 | 0 | 0 | 0 | 0 | 0 | 0 | 1 | 0 | 0 | 0 | 0 | 0 | 0 | 0 |
| Grade | 0 | 0 | 0 | 0 | 1 | 0 | 0 | 0 | 0 | 0 | 0 | | 0 | 1 | 0 | 0 | 0 | 0 | 0 | 0 | 0 | 0 | 1 | 0 | 0 | 0 | 0 | 0 | 0 | 0 |
| Predominant inflammatory cell type | 0 | 0 | 0 | 0 | N + M + L | 0 | 0 | 0 | 0 | 0 | 0 | | 0 | M + L | 0 | 0 | 0 | 0 | 0 | 0 | 0 | 0 | M + L | 0 | 0 | 0 | 0 | 0 | 0 | 0 |
| Peribronchial infiltrates | 0 | 0 | 0 | 1 | 0 | 1 | 1 | 0 | 1 | 0 | 0 | | 0 | 1 | 1 | 1 | 1 | 1 | 3 | 1 | 1 | 1 | 1 | 1 | 1 | 0 | 3 | 3 | 1 | 1 |
| Grade | 0 | 0 | 0 | 1 | 0 | 1 | 2 | 0 | 1 | 0 | 0 | | 0 | 1 | 1 | 1 | 1 | 1 | 3 | 1 | 1 | 1 | 2 | 1 | 1 | 0 | 2 | 2 | 1 | 1 |
| Predominant inflammatory cell type | 0 | 0 | 0 | M + L | 0 | M + L | M + L | 0 | M + L | 0 | 0 | | 0 | M + L | M + L | M + L | M + L | M + L | M + L | M + L | M + L | M + L | M + L | M + L | M + L | 0 | M + L | M + L | M + L | M + L |
| Necrotizing bronchitis | 0 | 0 | 0 | 0 | 0 | 0 | 0 | 0 | 0 | 0 | 0 | | 0 | 0 | 0 | 0 | 0 | 0 | 0 | 0 | 0 | 0 | 0 | 0 | 0 | 0 | 0 | 0 | 0 | 0 |
| Grade | 1 | 1 | 2 | 3 | 0 | 1 | 1 | 3 | 1 | 1 | 1 | | 1 | 0 | 1 | 1 | 2 | 1 | 1 | 1 | 1 | 1 | 1 | 1 | 1 | 1 | 1 | 1 | 1 | 1 |
| Predominant inflammatory cell type | M | M | M | M | 0 | M | M | M | M | M | M | | M | 1 | M + L | M | M | M | M | M | M | M | M | M | M | M | M | M | M | M |
| Intrabronchial mucus increase | 0 | 0 | 0 | 0 | 0 | 0 | 0 | 0 | 0 | 0 | 0 | not performed | 0 | M | 0 | 0 | 0 | 0 | 0 | 0 | 0 | 0 | 0 | 0 | 0 | 0 | 0 | 0 | 0 | 0 |
| Perivascular infiltrates | 0 | 0 | 0 | 1 | 0 | 1 | 1 | 1 | 0 | 0 | 0 | | 0 | 1 | 1 | 1 | 1 | 1 | 1 | 3 | 1 | 1 | 1 | 1 | 0 | 1 | 1 | 1 | 1 | 1 |
| Grade | 0 | 0 | 0 | 1 | 0 | 2 | 1 | 1 | 0 | 0 | 0 | | 0 | 1 | 1 | 1 | 1 | 1 | 1 | 1 | 1 | 1 | 1 | 1 | 0 | 1 | 1 | 2 | 1 | 1 |
| Predominant inflammatory cell type | 0 | 0 | 0 | M + L | 0 | M + L | M + L | M + L | 0 | 0 | 0 | | 0 | M + L | M + L | M + L | M + L | M + L | M + L | M + L | M + L | M + L | M + L | M + L | 0 | M + L | M + L | M + L | M + L | M + L |
| **Vascular lesions** | | | | | | | | | | | | | | | | | | | | | | | | | | | | | | |
| Intravascular rolling inflammatory cells | 0 | 0 | 0 | 0 | 0 | 0 | 1 | 1 | 0 | 0 | 1 | | 0 | 0 | 1 | 1 | 1 | 0 | 1 | 1 | 1 | 0 | 0 | 1 | 1 | 1 | 1 | 3 | 1 | 1 |
| Endotheliitis | 0 | 0 | 0 | 0 | 0 | 0 | 0 | 0 | 0 | 0 | 0 | | 0 | 1 | 0 | 0 | 0 | 0 | 0 | 0 | 0 | 0 | 0 | 0 | 0 | 0 | 0 | 3 | 1 | 0 |
| Vasculitis | 0 | 0 | 0 | 0 | 0 | 0 | 0 | 0 | 0 | 0 | 0 | | 0 | 0 | 0 | 0 | 0 | 0 | 0 | 0 | 0 | 0 | 0 | 0 | 0 | 0 | 0 | 1 | 0 | 0 |
| Hyaline thrombi | 0 | 0 | 0 | 0 | 0 | 0 | 0 | 0 | 0 | 0 | 0 | | 0 | 0 | 0 | 0 | 0 | 0 | 0 | 0 | 0 | 0 | 0 | 0 | 0 | 0 | 0 | 0 | 0 | 0 |
| Hemosiderin | 0 | 0 | 0 | 0 | 0 | 0 | 0 | 0 | 0 | 0 | 0 | | 0 | 0 | 0 | 0 | 0 | 0 | 0 | 0 | 0 | 0 | 0 | 0 | 0 | 0 | 0 | 0 | 0 | 0 |
| Arterial media hypertrophy/ hyperplasia | 0 | 0 | 0 | 0 | 0 | 0 | 0 | 0 | 0 | 0 | 0 | | 0 | 0 | 0 | 0 | 0 | 0 | 0 | 0 | 0 | 0 | 0 | 0 | 0 | 0 | 0 | 0 | 0 | 0 |
| Vascular occlusion | 0 | 0 | 0 | 0 | 0 | 0 | 0 | 0 | 0 | 0 | 0 | | 0 | 0 | 0 | 1 | 1 | 0 | 0 | 0 | 0 | 0 | 0 | 0 | 0 | 0 | 0 | 0 | 0 | 0 |
| **Diffuse alveolar damage (DAD)/Necrosis** | | | | | | | | | | | | | | | | | | | | | | | | | | | | | | |
| Necrosis alveolar epithelial cells (AEC) | 0 | 0 | 0 | 0 | 0 | 0 | 0 | 0 | 0 | 0 | 0 | | 0 | 0 | 0 | 0 | 0 | 0 | 0 | 0 | 0 | 0 | 0 | 0 | 1 | 0 | 0 | 0 | 0 | 0 |
| DAD | 0 | 0 | 0 | 0 | 0 | 0 | 0 | 0 | 0 | 0 | 0 | | 0 | 0 | 0 | 0 | 0 | 0 | 0 | 0 | 0 | 0 | 0 | 0 | 0 | 0 | 0 | 0 | 0 | 0 |
| Intraalveolar fibrin | 0 | 0 | 0 | 0 | 0 | 0 | 0 | 0 | 0 | 0 | 0 | | 0 | 0 | 0 | 0 | 0 | 0 | 0 | 0 | 0 | 0 | 0 | 0 | 0 | 0 | 0 | 0 | 0 | 0 |
| **Varia** | | | | | | | | | | | | | | | | | | | | | | | | | | | | | | |
| Hyperplasia/ hypertrophy bronchiolar epithelium | 0 | 0 | 0 | 0 | 0 | 0 | 0 | 0 | 0 | 0 | 0 | | 0 | 0 | 0 | 0 | 0 | 0 | 0 | 0 | 0 | 0 | 0 | 0 | 0 | 0 | 0 | 0 | 0 | 0 |
| Hyperplasia / hypertrophy type II pneumocytes | 0 | 0 | 0 | 0 | 0 | 0 | 0 | 0 | 0 | 0 | 0 | | 0 | 0 | 0 | 0 | 0 | 0 | 0 | 0 | 0 | 0 | 0 | 0 | 0 | 0 | 0 | 0 | 0 | 0 |
| Atypical cells / syncytia | 0 | 0 | 0 | 0 | 0 | 0 | 0 | 0 | 0 | 0 | 0 | | 0 | 0 | 0 | 0 | 0 | 0 | 1 | 0 | 1 | 0 | 0 | 0 | 0 | 0 | 1 | 1 | 1 | 0 |
| Pleura | 0 | 0 | 0 | 0 | 0 | 0 | 0 | 0 | 0 | 0 | 0 | | 0 | 0 | 0 | 0 | 0 | 0 | 0 | 0 | 0 | 0 | 0 | 0 | 0 | 0 | 0 | 0 | 0 | 0 |
| BALT | 0 | 0 | 0 | 0 | 0 | 0 | 0 | 0 | 0 | 0 | 0 | | 0 | 0 | 0 | 0 | 0 | 0 | 0 | 0 | 0 | 0 | 0 | 0 | 0 | 0 | 0 | 0 | 0 | 0 |
| TOTAL SCORE | 1 | 1 | 2 | 7 | 3 | 6 | 7 | 6 | 3 | 1 | 2 | - | 1 | 8 | 6 | 7 | 8 | 5 | 11 | 8 | 7 | 5 | 8 | 6 | 5 | 7 | 10 | 17 | 8 | 6 |
