## Supplementary Information Table 2 for "Enhanced fitness of SARS-CoV-2 variant of concern B.1.1.7, but not B.1.351, in animal models"

Histopathology score criteria developped by Angele Breithaupt, DipECVP

| **Overview** |
|---|
| **Intra-alveolar erythrocytes** (% affected area/lobe) (**NOT included in the score for mice)** |
| **Alveolar edema** (% affected area/lobe) |
| **Emphysema** (% affected area/lobe) **(NOT artefact)** |
| **Atelectasis** (% affected area/lobe) **(NOT artefact)** |
| **Higher magnification** |
| **Inflammation score** |
| **Alveolar infiltrates** (% affected area/lobe) |
| **Grade** 1 = single inflammatory cells; 2 = few inflammatory cells; 3 = almost complete atelectasis; 4 = complete atelectasis **(give representative grade for lobe)** |
| **Predominant inflammatory cell type:** Lymphocytes (L), Neutrophils (N), Macrophages (M), Plasma cells (P), Eosinophils (E), Mix |
| **Interstitial infiltrates** (% affected area/lobe) |
| **Grade** 1 = subtle infiltration; 2 = thickened walls but regular alveolar pattern; 3 = partially obscured architecture with thicker alveolar bridges; 4 = full atelectasis (**give representative grade for lobe)** |
| **Predominant inflammatory cell type:** L, N, M, P, E, Mix |
| **Peribronchial infiltrates** (including glands) (presence versus significant presence) |
| **Grade** 1 = 1 inflammatory cell layer; 2 = 2-3 infllammatory cell layers; 3 = 4-5 inflammatory cell layers; 4 ≥ 6 inflammatory cell layers (**give maximum grade)** |
| **Predominant inflammatory cell type:** L, N, M, P, E, Mix |
| **Necrotizing bronchitis** (presence versus significant presence) |
| **Grade** 1 = few inflammatory cells; 2 = inflammatory cell aggregates; 3 = inflammatory cells almost fill lumen; 4 > inflammatory cells completely fill lumen (**give maximum grade)** |
| **Predominant inflammatory cell type:** L, N, M, P, E, Mix |
| **Intrabronchial mucus increase: m**ain airways = 1; peripheral airways = 2 |
| **Perivascular infiltrates** (presence versus significant presence) |
| **Grade** 1 = 1 inflammatory cell layer; 2 = 2-3 inflammatory cell layers; 3 = 4 - 5 inflammatory cell layers; 4 ≥ 6 inflammatory cell layers (**give maximum grade)** |
| **Predominant inflammatory cell type:** L, N, M, P, E, Mix |
| **Vascular lesions** |
| **Intravascular rolling inflammatory cells** (presence versus significant presence) |
| **Endotheliitis** (presence versus significant presence) |
| **Vasculitis** (intramural damage, transmigrating inflammatory cells) (presence versus significant presence) |
| **Hyaline thrombi:** present = 1 |
| **Hemosiderin:** present = 1 |
| **Arterial media hypertrophy/ hyperplasia:** present = 1 |
| **Vascular occlusion:** present = 1 |
| **Diffuse alveolar damage (DAD)/Necrosis** |
| **Necrosis alveolar epithelial cells (AEC)**, NO hyaline membranes (% area/lobe affected) |
| **DAD** (AEC necrosis, hyaline membrane formation, debris, fibrin) (% area/lobe affected) |
| **Intraalveolar fibrin** present = 1 |
| **Varia** |
| **Hyperplasia/ hypertrophy bronchiolar epithelium** (cytoplasmic basophilia, multinucleated cells) (presence versus significant presence) |
| **Hyperplasia / hypertrophy type II pneumocyte**s (% affected area/lobe) |
| **Atypical cells / syncytia:** present = 1 |
| **Pleura:** mesothelium hypertrophy (% affected area/lobe) |
| **BALT:** present: # Follicle; # Germinal Center (GC) |
| **TOTAL SCORE** |

**Key:**
**% affected area/lobe (1 = below 5%; 2 = 6-40%; 3 = 41-80%; 4 > 80%)**
**Severity ( grade 1 - 4 depending on number of inflammatory cell layers)**
**Presence (1 - 3 foci) versus significant presence ( 3 = > 3 foci)**
